# Enabling multiscale variation analysis with genome graphs

**DOI:** 10.1101/2021.02.03.429603

**Authors:** Brice Letcher, Martin Hunt, Zamin Iqbal

## Abstract

**Background:** Standard approaches to characterising genetic variation revolve around mapping reads to a reference genome and describing variants in terms of differences from the reference; this is based on the assumption that these differences will be small and provides a simple coordinate system. However this fails, and the coordinates break down, when there are diverged haplotypes at a locus (e.g. one haplotype contains a multi-kilobase deletion, a second contains a few SNPs, and a third is highly diverged with hundreds of SNPs). To handle these, we need to model genetic variation that occurs at different length-scales (SNPs to large structural variants) and that occurs on alternate backgrounds. We refer to these together as multiscale variation.

**Results:** We model the genome as a directed acyclic graph consisting of successive hierarchical subgraphs (“sites”) that naturally incorporate multiscale variation, and introduce an algorithm for genotyping, implemented in the software `gramtools`. This enables variant calling on different sequence backgrounds. In addition to producing regular VCF files, we introduce a JSON file format based on VCF, which records variant site relationships and alternate sequence backgrounds.

We show two applications. First, we benchmark gramtools against existing state-of-the-art methods in joint-genotyping *17 M. tuberculosis* samples at long deletions and the overlapping small variants that segregate in a cohort of 1,017 genomes. Second, in 706 African and SE Asian *P. falciparum* genomes, we analyse a dimorphic surface antigen gene which possesses variation on two diverged backgrounds which appeared to not recombine. This generates the first map of variation on both backgrounds, revealing patterns of recombination that were previously unknown.

**Conclusions:** We need new approaches to be able to jointly analyse SNP and structural variation in cohorts, and even more to handle variants on different genetic backgrounds. We have demonstrated that by modelling with a directed, acyclic and locally hierarchical genome graph, we can apply new algorithms to accurately genotype dense variation at multiple scales. We also propose a generalisation of VCF for accessing multiscale variation in genome graphs, which we hope will be of wide utility.

## Background

Variant calling, the detection of genetic variants from sequence data, is a fundamental process on which many other analyses rely. There are two standard approaches, each with their own limitations. For Illumina data, mapping to a reference genome causes reference biases that affect discovery and genotyping: mapped reads favour the reference allele and reads in divergent regions fail to map [1–4]. For PacBio/Oxford Nanopore Technology (ONT) data, genomes can be fully assembled, and therefore the discovery and genotyping problems are in principle partially solved, by aligning each assembly against a reference. (There are caveats about how to get high per-base quality, either by hybrid ONT/Illumina or PacBio Hifi reads, but we leave this aside). However, the problem of how to coherently represent all of the variants in a cohort, comparing all against all, remains challenging both algorithmically and in terms of outputting results.

There are data structures that in principle can genotype alternate alleles which include both long structural variants and SNPs -some implementations include `Cortex`, `GraphTyper`, `vg`, `BayesTyper` [4–7]. All of these are based on graph representations of one form or another ranging from genotyping a whole-genome de Bruijn graph (`Cortex`), mapping all reads to a whole-genome Directed Acyclic Graph (DAG) of informative k-mers (`BayesTyper`), mapping all reads to a wholegenome graph of minimizing k-mers and matched haplotype index (`vg`/`Giraffe` [8]) or remapping premapped reads either to local DAGs of SNPs and indels off the reference (`GraphTyper`), or to graphs built from structural variant breakpoints (`GraphTyper2` [9]). These all reduce the impact of reference bias, and allow cohort genotyping at consistent sites, but all of them struggle with the issue of representation. We highlight two important types of situations which pose representation challenges -any good solution will need to address these, and conversely, solving these alone would address most of the needs of most users.

First, a long deletion that “covers” 10 SNPs will in principle have 2^10^ alternate alleles. This would be painful to output in the widely-used Variant Call Format [10] (VCF), but more fundamentally, it would force the genotyper to make statements about long alternate alleles that only differ by SNPs. If two long alleles, one true and one not, differed by just one SNP, then under most models they would have very similar likelihoods, and it would be impossible to tell clearly which was the correct allele. In other words, there would be no fine-scale data about variants, making SNP filtering impossible (Supplementary Fig. 1).

Indeed, there is no tool (to our knowledge) that self-advertises as supporting joint structural variant and SNP genotyping. Benchmarking in this paper shows `GraphTyper2` and `vg` both work in this scenario if given VCF input. The second key challenge occurs when variants occur on different genetic backgrounds such as diverged MHC haplotypes [3] or large insertions -for this there is no current solution.

We highlighted above two key situations to address: SNPs as alternatives to long deletions, and SNPs on top of long alternate haplotypes. In both of these cases variants are bound by relationships. In the first, they are mutually exclusive and in the second, they occur on top of alternative sequence backgrounds, thus combining exclusion and a hierarchy. We call such variation **nested**, and identify these relations as sufficient to capture a valuable proportion of natural genetic variation. We therefore model the genome as a directed acyclic graph that is a succession of locally hierarchical subgraphs. This is a rich enough model to incorporate our key use-cases, without incurring the price of excessive generality; we discuss the benefits and limitations in the Discussion. Based on this, we can use `gramtools` to identify these nested site relationships, leverage them during genotyping and output variants, genotypes and likelihoods in a file format extending VCF. This is, to our knowledge, the first framework for jointly analysing genetic variation at different scales (SNPs and structural variants) and on different sequence backgrounds.

We start by detailing the genome graph workflow implemented in `gramtools` and an algorithm for genotyping nested variation. We build graphs of variation from 2,498 samples at four *Plasmodium falciparum* surface antigen genes which harbour high levels of diversity, including two that each have two diverged allelic forms (DBLMSP and DBLMSP2). We use simulated haplotypes from graphs to evaluate the impact of nesting on genotype confidences, and then benchmark with empirical data from 14 *P. falciparum* samples which have high quality PacBio assemblies for ground truth. We then address the two canonical use-cases we highlighted above. We show that `gramtools` outperforms `vg` and `GraphTyper2` when genotyping long deletions and all the overlapping small variants from a cohort of 1,017 *Mycobacterium tuberculosis* genomes. Finally, we apply `gramtools` to the use-case for which no current solution exists. We genotype 706 African and SE Asian *P. falciparum* genomes at the gene DBLMSP2, which possesses variation on two diverged backgrounds which had previously appeared to either never, or very rarely, recombine. This generates the first map of genetic variation on both diverged backgrounds, revealing patterns of recombination that were previously unknown.

## Results

`gramtools` implements a workflow for building, genotyping and augmenting genome graphs (Fig. 1). Genotyping serves two main use cases in this workflow. First, it is used for inferring a sample’s closest path in the graph, which we can use as a “personalised” reference genome, since it should be a closer approximation than any individual genome would be. We are thereby able to discover new variants by using standard reference-based variant callers with this personalised reference, an approach previously described in [3, 11, 12]. Second, `gramtools` is used for genotyping cohorts of samples on a graph containing all such discovered variants. Neither case requires finding variants absent from the graph because novel variants are found by standard tools applied to the personalised reference. Thus while other tools such as `vg` [4] perform full alignment of reads to the graph, `gramtools` achieves the same aims while only needing to do exact matching. Our approach is currently limited to high-accuracy short reads (e.g. Illumina); support for long erroneous reads requires matching read substrings.

**Figure 1.**
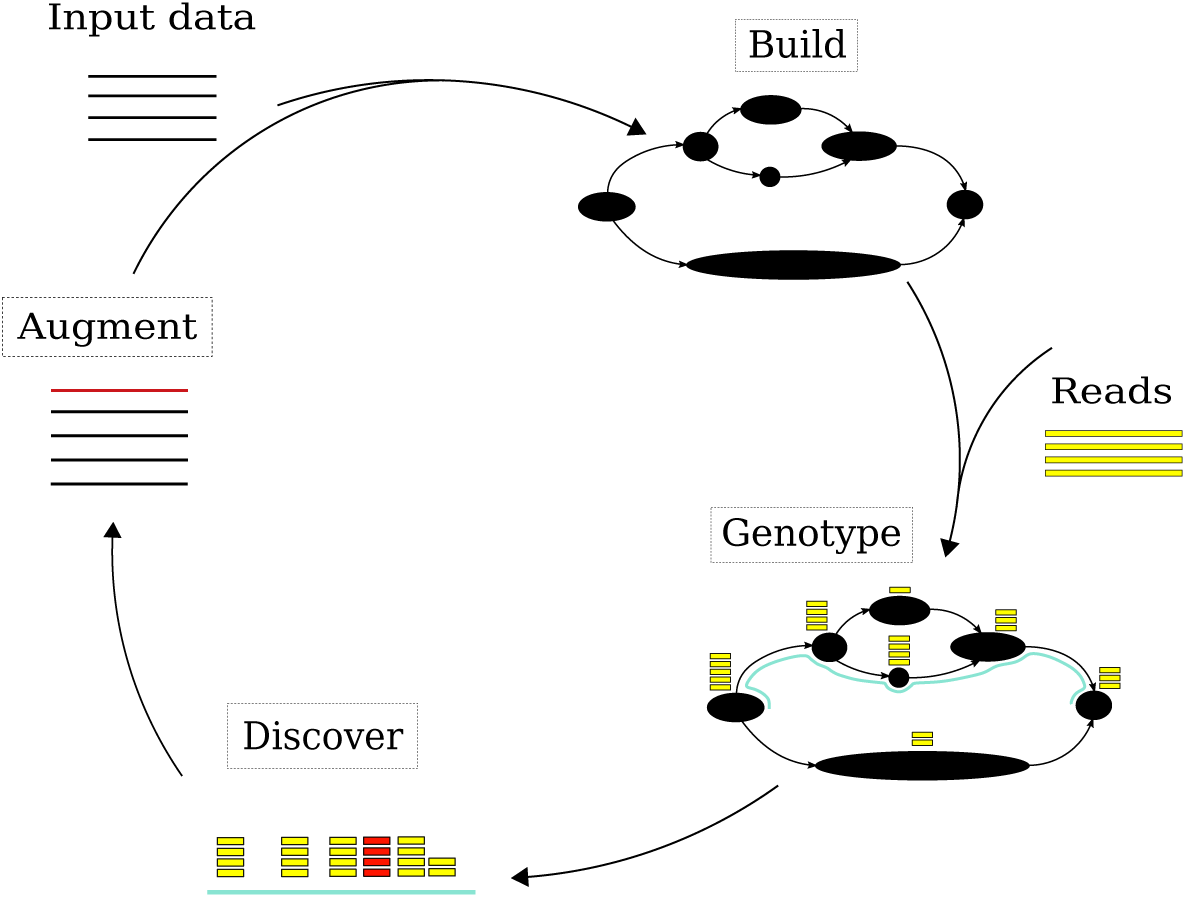
Genome graph workflow implemented in gramtools. Black nodes represent genomic sequence or site entry/exit points. Build consists in producing a genome graph that is directed and acyclic, a requirement for gramtools. Genotyping consists in calling alleles at each variant site and inferring a haploid personalised reference genome for a sample. New variants, shown in red, are discovered using standard reference-based callers run against the personalised reference.

### Graph constraints and genotyping with the vBWT

In `gramtools` sequence search in genome graphs is supported using the compressed suffix array [13] of a linearised representation of the graph, which we call variation-aware Burrows-Wheeler Transform (vBWT). The vBWT was developed in [11]; details of how it converts BWT string matching to graph mapping are provided in the Methods. A key requirement of the vBWT is that the graph be decomposable into a succession of subgraphs (sites) each of which is strictly nested (see Fig. 2 and Methods for formal definitions) interspersed by linear non-variable regions.

**Figure 2.**
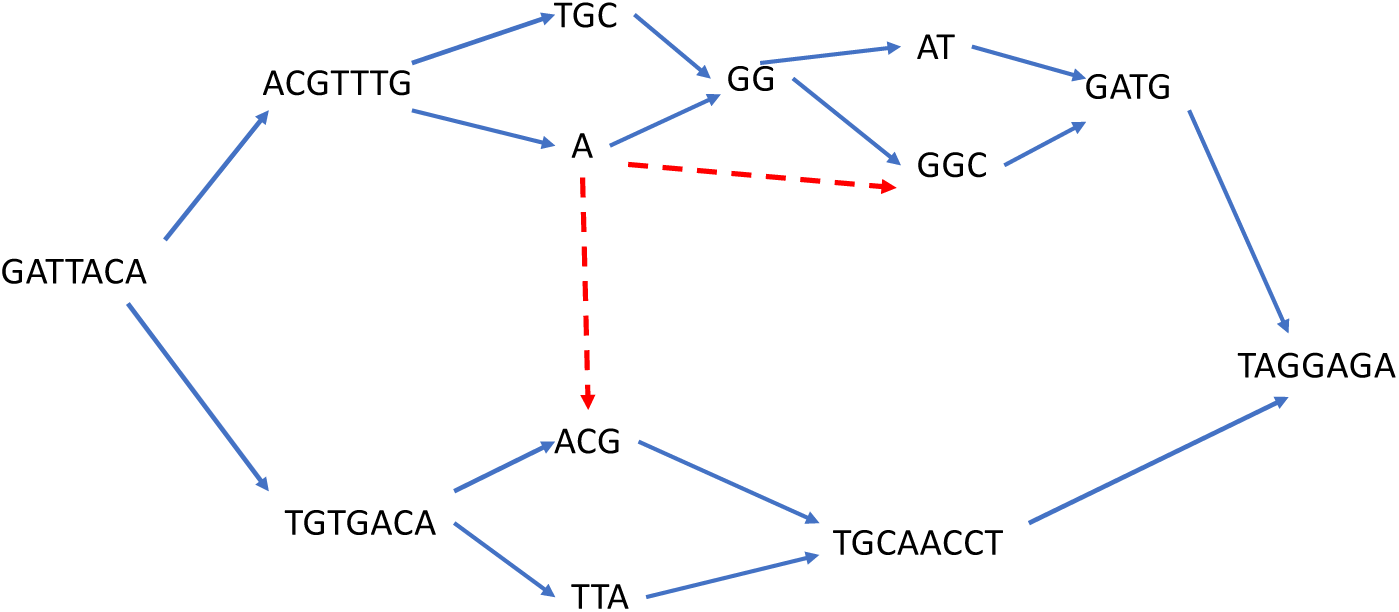
Gramtools requires variation to be expressed as a nested directed, acyclic graph (DAG). A nested DAG represents the genome as a DAG with a single source and sink, which can be decomposed into a succession of subgraphs (or sites). Each site starts with an opening node, and finishes with a closing node, and consists of strictly nested sub-sites. This allows hierarchical genotyping of alternate alleles. Strict nesting means that sub-sites must close off and complete before their parent site, and without connecting to different sub-sites (e.g. the dotted red lines would not be permitted).

This results in a graph where genotyping a site with alternate alleles is well-defined, preserving a notion with biological value, but also places some restrictions on the structure of the genome graph. We give an example in Supplementary Fig. 2 showing a pair of allowed/disallowed graphs that generate equivalent sequence.

These graphs can be built from multiple sequence alignments (MSAs) or a reference genome plus VCF file; we used MSAs in this paper. The construction process, first introduced in [14], is explained further in the Methods and we consider the implications of this model in the Discussion. The original vBWT did not support nesting at all, and in this paper we introduce our nesting implementation; in addition we have optimised the codebase to improve mapping, coverage recording, and genotyping.

### Genotyping nested genome graphs

`gramtools` genotypes a nested DAG in which variant sites have been defined (see Methods). `gramtools` genotypes individual sites by choosing the maximum-likelihood allele under a coverage model that draws on ideas from `kallisto` [15], including both per-base coverage information, and equivalence class counts for reads that could equally support different alleles (see Methods). Genotype likelihoods are calculated for each allele at a site, and the ratio of likelihoods between the maximum likelihood allele and the next best is termed the genotype confidence. With nested variation, we apply this model recursively from child sites to their parents, with candidate alleles in parent sites generated based on the genotype calls of child sites (see Methods).

An example of the nested genotyping procedure is shown in Fig. 3. To maintain coherence, if child sites on two different branches of a parent site are genotyped, whole branches can get invalidated. For example at a ploidy of one if an outgoing branch from a parent site is called, all children sites on the other branches get null calls. We refer to each outgoing branch from a parent site as a **haplogroup**, for a group of related haplotypes.

**Figure 3.**
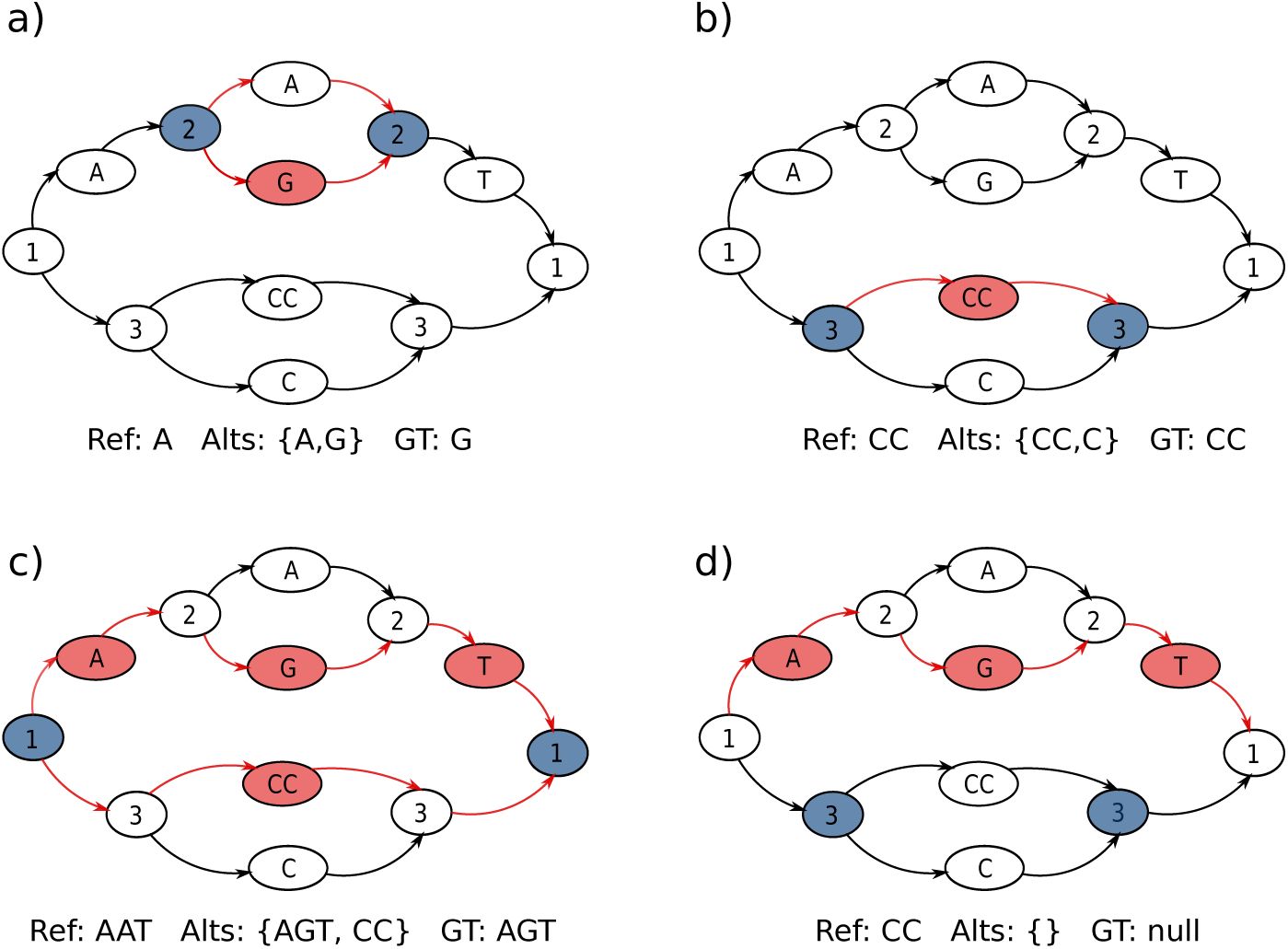
Nested genotyping procedure. Nodes with numbers mark variant sites. In each panel, blue-filled nodes mark which site is being processed, red-filled nodes mark called alleles, and red paths mark alleles considered for genotyping. *Ref* is the reference allele, *Alts* are the alleles considered for genotyping, and *GT* is the called genotype. The example shows haploid genotyping. Genotyping of child site 2. **b)** Genotyping of child site 3. **c)** Genotyping of parent site 1, with *Alts* generated from the called alleles in child sites 2 and 3. **d)** The model genotypes the path going through site 2, causing site 3 to be invalidated (null called).

### jVCF output format

Nested graphs create two problems for outputting genotyped sites. First, the genomic positions of output records can overlap, implying they should be considered jointly. Second, sites can occur on sequence backgrounds (haplogroups) that do not include the reference genome sequence. In VCF, overlapping records require careful genotyping, and alternate references are not supported.

Storing the parent/child relationships between sites in nested graphs provides a clear solution. For the first problem, it makes incompatibilities between sites explicit, allowing genotyping to enforce consistency (Fig. 3). For the second problem, it enables defining haplogroups and which ones variants fall on. This in turn allows defining alternate reference sequences, coordinates and names.

`gramtools` outputs a variant call file in JavaScript Object Notation (JSON), a widely used format for storing data as key-value pairs. The file stores variant records mirroring VCF and additionally stores parent/child site relationships -we thus call it jVCF. By storing site relationships, jVCF provides a language to refer to variant sites in terms of other sites and of haplogroup background, allowing the definition of alternate references and queries such as extracting all variant records under a given haplogroup. A format specification and toy example are provided in the Supplementary text. In addition to a jVCF file, `gramtools` outputs a regular VCF file containing only non-nested sites, yielding a VCF file with no overlapping records and referring only to the linear reference genome.

### Validation of nested genotyping with simulated data

Our first simulation was designed to evaluate both genotyping performance, and whether nested genome graphs resulted in improved calibration of genotype confidences. We based the simulation on a real example where there are two alternate haplotypes each bearing variants, building graphs of *P. falciparum* variation for two genes, DBLMSP and DBLMSP2. These genes exhibit a dimorphism at the Duffy Binding-Like (DBL) domain, each having a region *>*500bp in length with two allelic forms that are highly diverged [16, 17] (around 65% sequence similarity [18]). We built two versions of the graph, one without any nesting, and one allowing nesting up to five levels deep (see Methods). The graphs were built from high confidence variant calls in 2,498 samples from the Pf3k project [19] (see Methods for details). The graph without nesting contained 451/413 variant sites for DBLMSP/DBLMSP2 respectively, and the graph with nesting contained 558/500 variant sites respectively, as nesting allowed SNPs/indels on alternate haplotypes to become sites in their own right. We randomly sampled 10 paths from the non-nested graph (which therefore exist in the nested one), recorded the implied truth variant calls, simulated reads from the paths, and passed them to `gramtools` for genotyping (see Methods for details). Out of 17,280 evaluated calls in the non-nested graph, gramtools recovered 99.9% (recall) and of all calls made, 99.8% were correct (precision). In the nested graph, out of 21,160 evaluated calls, recall and precision were both 99.9%.

While `gramtools` genotypes both types of graphs to high accuracy, nested graphs provide better call resolution. This is shown in Fig. 4 for DBLMSP2 where the nested graph reflects the allelic dimorphism: SNPs/small variants fall on top of each allelic form. We also confirm that the genotype confidence of correct calls is also increased, since nesting allows likelihood calculations based on coverage precisely at SNPs on alternate haplotypes, rather than an average across the whole haplotype (coverage shown in Figure. 4, and effect on confidences shown in Supplementary Fig.4).

**Figure 4.**
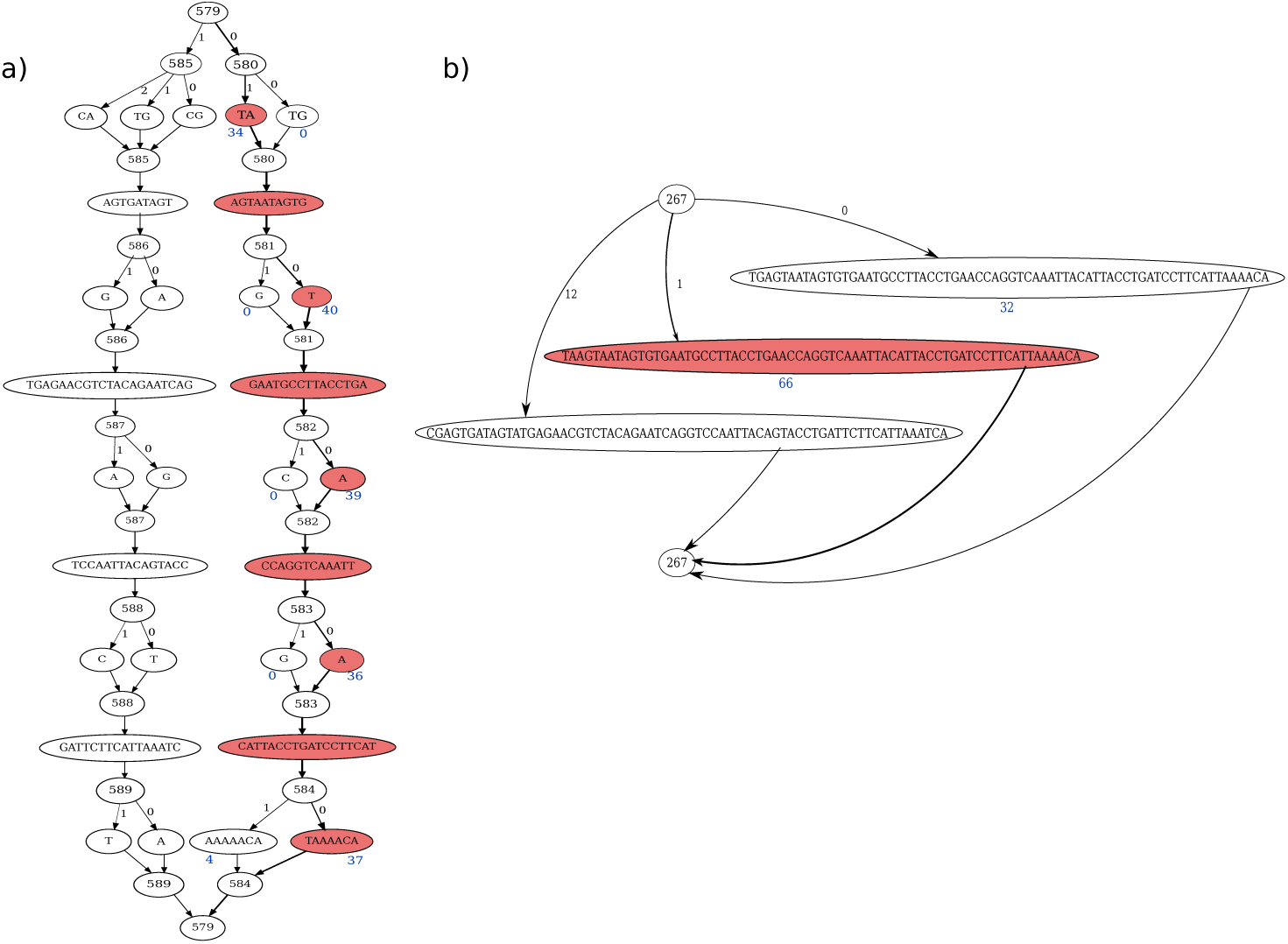
Nested graphs improve site resolution and coverage differences. A call for part of DBLMSP2 is shown (*P. falciparum* reference genome Pf3D7 chromosome 10 positions 1433921:1433987). Red nodes mark the called allele and spell the same sequence across nested (panel a) and non-nested (panel b) graphs. Numbered nodes mark variant sites and edges are labeled by haplogroup. Blue text under the nodes gives read coverage for the called and next best allele. In the non-nested graph the next best allele is long and only one SNP away from the best allele so that reads mapping to common sequence add coverage to both. This reduces coverage differences compared to the nested graph. 3 out of the 13 alleles are shown in panel b for clarity.

### Empirical evaluation against single-reference tools at surface antigens

We set out to evaluate the `gramtools` workflow in comparison with standard single-reference callers `SAMtools` [20] (the classical “pileup” variant caller) and `Cortex` [5] (which discovers bubbles in a de Bruijn graph, and then maps flanking sequence to the reference to get coordinates). We include `Cortex` because it has previously been shown to produce high quality calls even in the indel-rich *P. falciparum* genome, and can successfully identify alternate alleles at the genes we analyse here [21], but has no capacity to consider nested variation. We built a whole-genome graph containing variation from 2,498 *P. falciparum* samples (as in the simulation experiment) in four surface antigen genes: DBLMSP, DBLMSP2, EBA175 and AMA1 (see Methods). The additional genes, EBA175 and AMA1, are both vaccine targets, where there is great value in being able to correctly identify known and novel variation [22, 23]. We used 14 *P. falciparum* validation samples with both Illumina data and high quality PacBio long-read assemblies [24], and which had been excluded from graph building, to assess `gramtools` genotyping. `SAMtools` and `Cortex` were each run twice, once with each of two reference genomes. The first was the standard *P. falciparum* 3D7 reference genome [25]. The second was the closest haploid path in the graph to the sample, i.e. the personalised reference (PR) genome described above. The PR allows reference-based variant calling in a genome graph context [11, 12] and is produced by `gramtools` during genotyping (see Methods). Performance is measured as the edit distance between the gene sequence with called variants applied and the long-read assembly, normalised by gene length.

We show in Fig. 5 the scaled edit distance achieved by these tools on the 14 validation samples in DBLMSP2, as compared with just taking the 3D7 (reference) allele.

**Figure 5.**
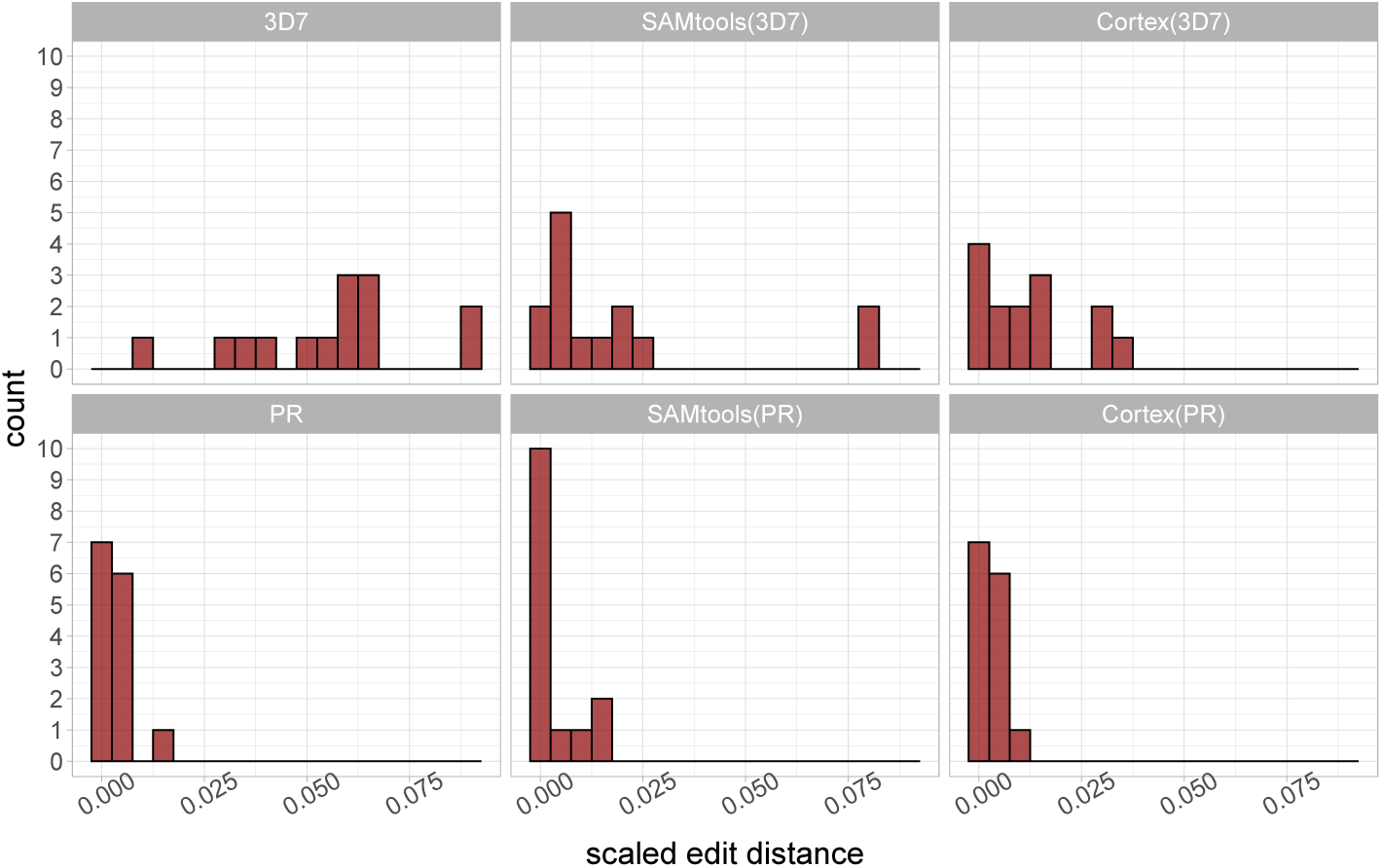
Histogram of accuracy of different tools in inferring sequence of DBLMSP2 gene in 14 validation samples. In all panels, x-axis is the scaled edit distance (i.e. edit distance of inferred gene sequence from the true sequence determined by high quality PacBio assembly, divided by length of DBLMSP2), and y-axis is the number of samples in that bin. The two histograms in the left-hand column show how close the 3D7 reference genome, and the personalised reference (PR) are to each sample. The two histograms in the middle column show results for SAMtools when using the 3D7 and PR as reference genome. The histograms in the right-hand column are the equivalent plots for Cortex. In all cases, the PR is a clear improvement on the standard reference. Mean scaled edit distances are 5.52%, 1.90%, 1.25%, 0.37%, 0.35%, 0.32% for 3D7, SAMtools (3D7), Cortex (3D7), PR, SAMtools (PR) and Cortex (PR) respectively.

The standard inference approach is to map to the 3D7 reference and use a pileup-based approach to variant calling, such as `SAMtools`. Since DBLMSP2 has two deeply diverged haplotypes, any sample with the non-3D7 haplotype does badly with this approach, and we see `SAMtools` achieves a scaled edit distance of over 7.5% for 2/14 samples. `Cortex`, which uses assembly, improves on this, but still leaves 3 samples with over 2.5%. However, looking at the bottom 3 histograms, we see the results based on the `gramtools`-inferred PR all improve on these results. `SAMtools` achieves more perfect results (10/14) and `Cortex` has more samples within 1% scaled edit distance (13/14). These results validate the use of a graph over a single reference for these diverse genes, and also confirm the benefit of using `SAMtools` and `Cortex` with the gramtools-inferred PR, showing that the PR provides a bridge between the standard reference and the truth.

Results for DBLMSP, AMA1 and EBA175 are broadly similar; `gramtools` genotyping outperforms reference-based callers using 3D7 (Supplementary Figures 5,6,7). The results are dramatic for EBA175, as the graph contains the two dominant alleles (one long, one short), so even the PR itself without `SAMtools`/`Cortex` is near perfect. AMA1 is similar (the PR has edit distance 0 for 13/14). In DBLMSP four samples remain at a distance of 4-10% to the truth assemblies. These appear to be genomes containing variation neither captured in the graph nor callable by `SAMtools` and `Cortex`, and would be natural candidates for long-read sequencing and then incorporating their alleles into the graph.

#### Comparing gramtools personalised reference with best of reference panel

The first step of the `gramtools` workflow involves inferring a PR, a recombinant of the panel of genomes used to construct the genome graph. We measured, on DBLMSP2 for the 14 validation samples, the benefit of using this PR compared with just choosing the closest sequence from the input panel (see Methods for how the closest sequence is determined). We show the results in Fig. 6, plotting the difference between two edit distances -the distance of the PR from the truth, and the distance of the best panel reference and the truth (where truth refers to the sequence in the PacBio assembly of the relevant sample). The sequences with negative values indicate recombinants found by `gramtools` that are closer to the truth assembly than any input to the graph. Thus for 9/14 samples the PR was superior to best-input, and for 3/14 they were equally good. Similarly for AMA1 the PR was as good or closer to the truth 13/14 times, (Supplementary Fig. 8). EBA175 is dominated by single biallelic indel, both alleles of which were in the input data, so the PR was identical to the best input in 11/14 cases, and better in 2/14 (Supplementary Fig. 9). Finally, for DBLMSP, there were 2 samples for which the PR was 5-6% more distant than the best input in the graph (Supplementary Fig. 10). We found this is due to a specific unresolved region where sites had null calls due to low coverage.

**Figure 6.**
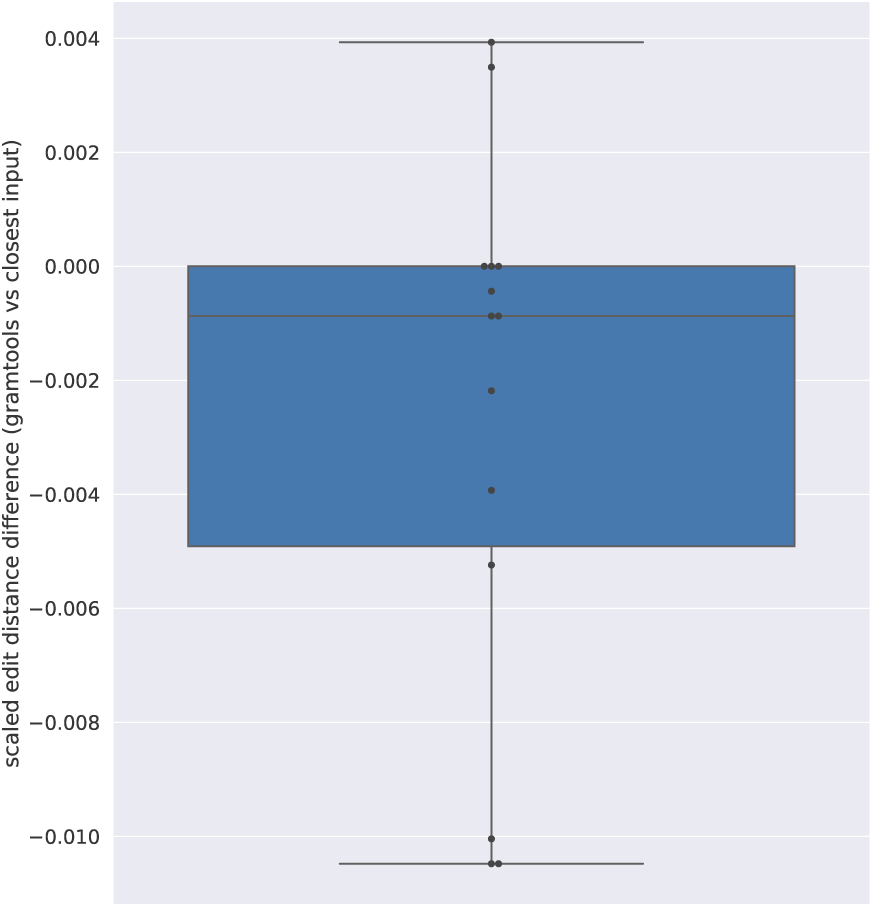
gramtools genotyping finds recombinants between input haplotypes in the graph. Each dot shows the difference between the distance of the gramtools-inferred sequence from the truth, and distance of the closest input sequence to the truth. Positive values indicate that one of the input sequences is closer to the truth than the *gramtools*-inferred sequence, while negative values indicate `gramtools` finding recombinants in the graph that are closer to the truth than any input sequence.

### Application: unified SNP and large deletion analysis in *M. tuberculosis*

Nested variation occurs naturally when jointly genotyping small variants overlapping structural variants. We performed an experiment to assess how `gramtools` compares to two other genome graph tools, `GraphTyper2` [9] and `vg` [4], in one such situation in *M. tuberculosis*. The experiment evaluates each tool’s ability to genotype a fixed set of input SNPs and overlapping deletions.

We started from variant calls obtained by running `Cortex` on 1,017 publicly available Illumina samples (see Methods). 17 of these samples have matched Illumina [26] and PacBio reads [27], from which we produced high quality hybrid assemblies (see Methods). These assemblies were used as ground truth for evaluating genotyping. We identified 73 high quality large deletion calls in the 17 samples, confirmed using the assemblies (see Methods). We then extracted all variation in the 1,017 samples overlapping these deletions. Together these provide the variant sites at which we evaluate each tool.

For `gramtools` we built a genome graph of each deletion region from multiple-sequence alignments of the `Cortex` variant calls applied to the *M. tuberculosis* H37Rv reference genome [28] (see Methods). The graphs were then combined with the rest of the reference genome. To genotype the same variants in `GraphTyper2` and `vg`, we merged the VCF files of all 1,017 input samples using `bcftools`. VCF is the required input format for genotyping in `GraphTyper2` and the only input format that worked in `vg` after failing on multiple sequence alignments (see Methods).

Altogether the deletion regions cover 51,701 bp of the reference genome. The variants under them cover 4,105 reference positions in 1,109 sites in the `gramtools` graph and 2,386 positions in 1,434 sites in the merged VCF file.

We first looked for each of the 73 known deletions in each tool’s VCF output and found `GraphTyper2` called all 73, `gramtools` called 70 and `vg` called 66. We then assessed each tool’s ability to resolve each deletion region in the 17 evaluation samples. For each region, we applied called variants to the *M. tuberculosis* reference genome and measured edit distance to the truth assembly using the `minimap2` [29] aligner (see Methods).

Fig. 7 shows the cumulative distribution of scaled edit distances (edit distance divided by the length of the sequence) for each tool. `gramtools` achieves the lowest mean distance to the truth (1.18%), followed by `GraphTyper2` (2.02%), `vg`(2.36%), and using the reference genome sequence alone (4.78%). Correspondingly, the fraction of perfectly resolved sequences (edit distance 0) is 86.7%, 66.8%, 54.0% and 43.3% respectively. Thus, while all tools perform better than using the reference sequence alone, `gramtools` shows improved ability to jointly resolve large deletions and small overlapping variants compared to state-of-the-art tools `vg` and `GraphTyper2` on this dataset.

**Figure 7.**
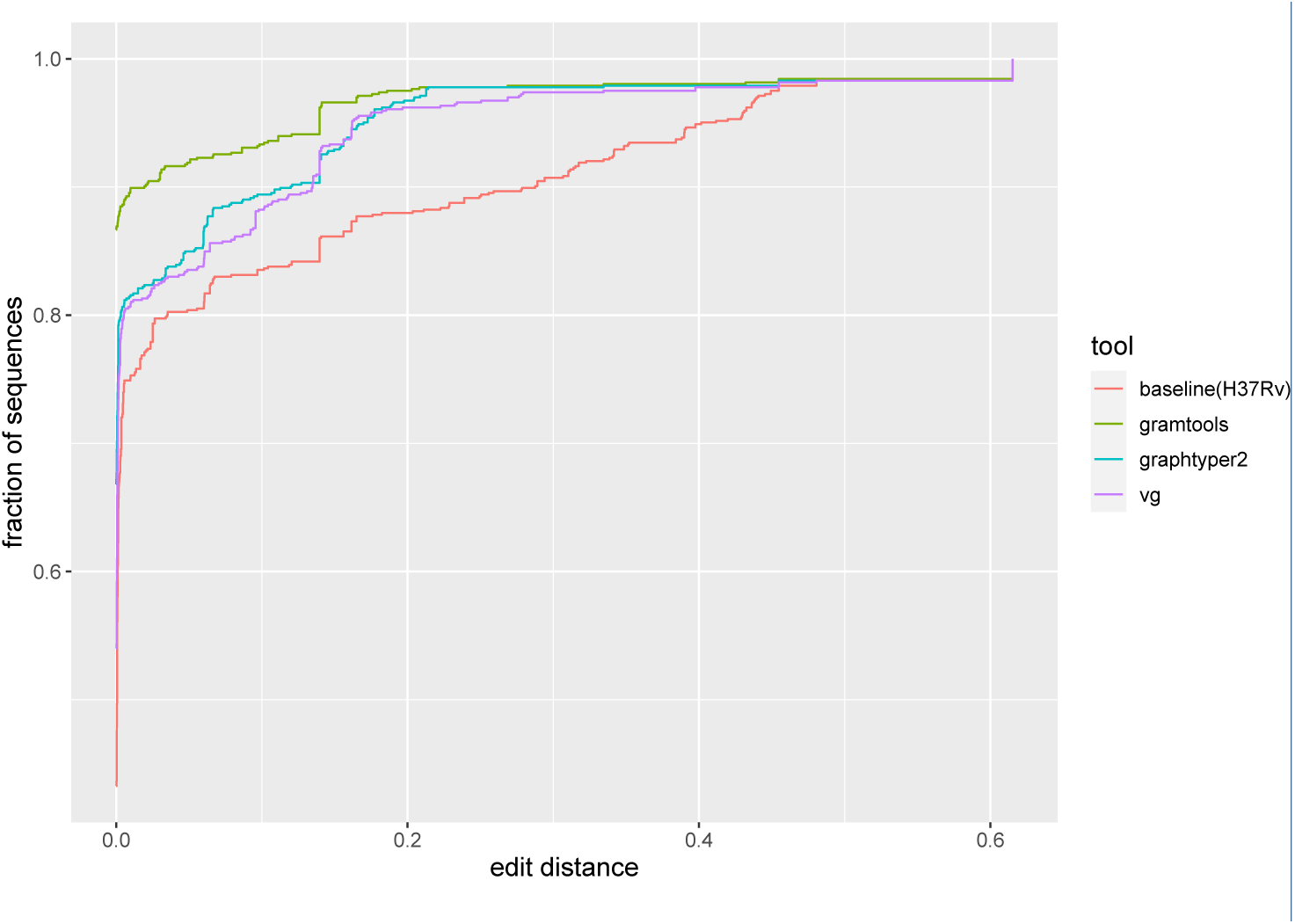
gramtools joint SNP and deletion genotyping performance compared to other genome graph tools. The curves are the cumulative frequencies of edit distance between inferred sequences and truth assemblies. *baseline* refers to using the *M. tuberculosis* H37Rv standard reference sequence only. `gramtools` achieves the lowest mean edit distance and the highest fraction of perfectly resolved sequences (edit distance 0).

For completeness, results using `bowtie2` [30] instead of `minimap2` alignments are shown in Supplementary Fig. 13 and give the same performance trends but fewer aligned sequences.

In terms of computational performance, `gramtools` processed the most reads per CPU second while using comparable amounts of RAM on this dataset (Table 1). A bottleneck in `vg` is temporary disk use, exceeding 500 Gigabytes without pruning the graph to remove densely clustered variation. For `GraphTyper2`, we include counting a separate mapping step to the reference genome (with `bowtie2`) as it is a prerequisite to genotyping.

**Table 1.** Computational performance of each tool. *Index*: vg genome graph built from VCF and pruned to reduce graph complexity (else temporary disk use exceeded 500 Gb before completion, see Methods). `GraphTyper2` runs from a VCF file and has no separate indexing operation. *Genotype*: Speed shows the average number of reads mapped across the 17 samples (10.7 million) divided by the average CPU time. vg and `GraphTyper2` have separate read mapping and genotyping steps: for speed, CPU time is summed, and for RAM, mapping is shown followed by genotyping in brackets. `GraphTyper2` does not implement its own mapping but requires an input file of reads mapped to a linear reference genome; mapping RAM and speed is shown for `bowtie2` with default parameters. *metrics*: Mb: Megabytes; sec: total CPU seconds (accounts for multi-threading, 10 threads used for genotyping in each tool).

|  | Index |  |  | Genotype |  |
| --- | --- | --- | --- | --- | --- |
|  | Disk(Mb) | Max RAM(Mb) | Speed(sec) | Max RAM(Mb) | Speed(reads/sec) |
| vg | 29 | 609 | 105 | 605(158) | 3,961 |
| gramtools | 153 | 480 | 32 | 632 | 34,290 |
| GraphTyper2 | - | - | - | 869(88) | 7,604 |

While performance of `gramtools` is better here -especially compared to `vg` on unpruned graphs -we note that genome graph tool performance depends in complicated ways on genome size, and repeats, and the amount and density of stored variation. We are for example aware of `gramtools`’ much lower mapping speed to the human genome [31] and therefore emphasise that different tools are likely tuned for specific use cases -some species like humans have large genomes with low diversity while others, like the examples in this paper, have small genomes but higher diversity.

In this experiment `GraphTyper2` and `vg` are able to genotype variation at multiple scales. One caveat is the VCF file they genotype contains inconsistencies. For example, if one VCF record describes a deletion and another describes an overlapping SNP, a reference call at the deletion and an alternate call at the SNP are inconsistent because the two calls imply different sequences. This occurs because the variants are related but expressed in isolation. By contrast `gramtools` models site relationships explicitly, outputting a VCF file without inconsistencies and a jVCF file mapping the nested variation.

An output format like our proposed jVCF becomes especially important when analysing more complex variation such as SNPs on top of alternate haplotypes, where variants need to be expressed against different references. We now show such an application of multiscale variation analysis using the *P. falciparum* surface antigen DBLMSP2, which would not be possible using the VCF files output by `vg` or `GraphTyper2`.

### Application: charting SNPs on top of alternate haplotypes

When two diverged forms of a gene segregate in a population, we want to access small variants on top of each. Returning to the surface antigen DBLMSP2 in *P. falciparum* -which we have shown is accurately genotyped by `gramtools` using simulated and real data -we assessed whether `gramtools`’ multiscale genotyping and jVCF output could recover the two diverged forms of DBLMSP2 and access variation on top of each form. We genotyped 706 *P. falciparum* samples from Ghana, Cambodia and Laos using `gramtools` and analysed a combined jVCF file of all calls in all samples (see Methods). Genotyping, including read mapping, used an average of 1.14 Gigabytes of peak RAM, processing an average of 2,525 reads per CPU second.

Fig. 8 shows a matrix of calls in each sample inside the DBL domain of DBLMSP2, known to be dimorphic [32]. Hierarchical clustering distinguishes two groups of samples above and below the left-hand side red arrow, with an average scaled edit distance between the two groups of 16.8% compared to within-group distances of 1.4% and 4.8%. `gramtools` can thus recover the two divergent forms.

**Figure 8.**
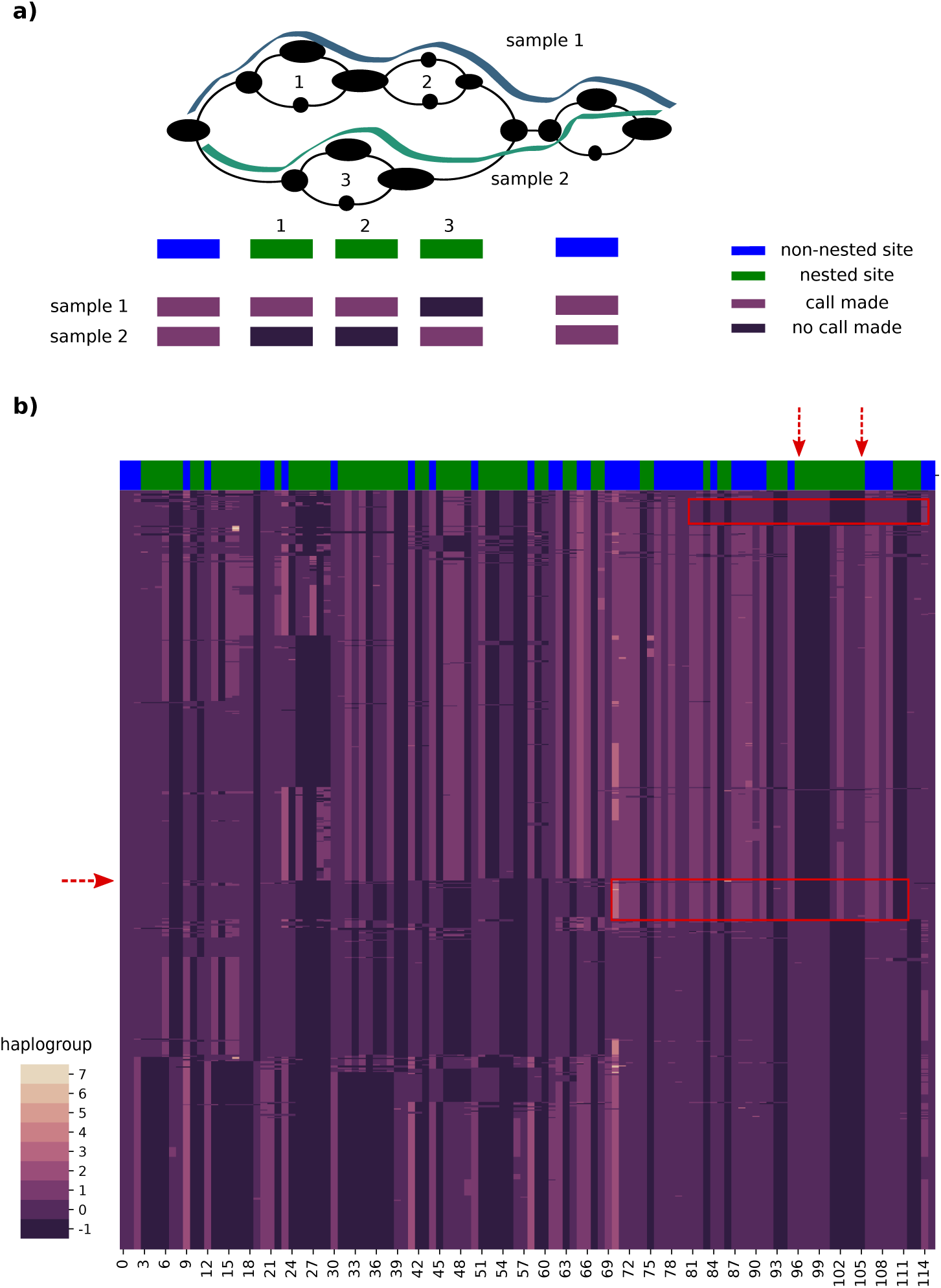
gramtools captures allelic dimorphism and nested variation in DBLMSP2. **a)** A toy example of a graph with SNPs on top of diverged haplotypes. Each column is a variant site laid out linearly by relative genomic position. The top row labels sites green if they are nested inside other sites and blue if not. The next two rows show two haploid genotyped samples: sites labeled 1 and 2 are genotyped mutually exclusively to site labeled 3. **b)** Each row is one of 706 samples and each column is a variant site. As in panel a, green and blue bars at the top denote nested and non-nested sites. A haplogroup of -1 indicates no call made at a site. Clustering of the samples divides them in two groups above and below the red arrow on the left, reflecting the two forms of the gene. The top red arrows enclose a set of nested SNPs on two divergent haplotypes: as in panel a, samples are genotyped mutually exclusively at these sites depending on which form of the gene they have. The red rectangles on the right-hand side highlight recombinant samples: the haplogroup patterns look like one gene form on the left of the rectangles, and like the other gene form inside the rectangles.

`gramtools` also produces calls in nested sites, shown as green bars at the top of the matrix. In between the two red arrows at the top, a block of nested sites is called mutually exclusively: when one of the gene forms has a genotype call, the other form receives a null call (shown as black-coloured cells), and vice-versa. The variants occur on different sequence backgrounds and thus get selectively invalidated (as per the algorithm in Fig.3). This demonstrates `gramtools`-defined haplogroups provide access to small variants on top of each form. It further shows the genome graph structure is reflecting the dimorphism. One consequence of this analysis is that it allows us, for the first time to our knowledge, to see clear evidence of recombination in the dimorphic region. We highlight in red boxes two regions where there is clear evidence of recombination, containing different haplogroup patterns than directly to the left of the boxes.

In this analysis the nested calls and their haplogroup background can only be accessed using `gramtools`’ jVCF output. Storing site relationships allows `gramtools` to enforce genotyping coherence and to define alternate reference backgrounds on which individual variant sites lie.

## Discussion

Genetic variation occurs through different mechanisms at different scales ranging from SNPs to large structural variants. The need to jointly analyse SNPs and structural variants therefore arises immediately on trying to genotype a cohort. We have presented a method for identifying, calling and outputting such variation in

`gramtools`. By identifying site relationships in the graph, `gramtools` is able to geno-type incompatible sites mutually exclusively and to output variation both against the standard reference genome and against locally defined alternate references.

One of the challenges of extending linear references with graphs is recognising modelling assumptions. Working from a single reference implicitly assumes that individuals within a species have genomes that are close to the reference. Where this model breaks down (divergent sequence, repeats, structural variation) artefacts appear and become common rules of thumb to check for: clustered heterozygous calls suggest a missing repeat, patterns in read pair orientation and spacing suggest structural variation [33]. When moving to a graph, we are forced to make new modelling choices.

At one end of the spectrum is the `gramtools` approach: genome graphs must be nested, directed acyclic graphs (NDAGs). This simple model allows direct access to two key notions we want to use: easily distinguishing horizontal (paralog-like) and vertical (ortholog-like) variation, and defining distinct alternate sequence backgrounds. At the other extreme are very general sequence graphs with no ordering, whether De Bruijn (which collapse all repeats of size *k*) or `vg`-like (bidirected and allowing cycles). These models better support more complex events such as duplications and inversions, with an added cost in complexity of implementation. We would argue it is unlikely a single model is best for all problems and one should choose the approach best suited to the biological and genetic question at hand. To that end, it would be valuable for the pangenomics field to develop standardised use-cases and evaluation datasets. This would help us avoid over-optimising for specific datasets, and help understand the different performance challenges of repetitiveness (*P. falciparum* is much more repetitive than human), genome size (microbes are tiny but diverse) and loci where alternate reference haplotypes are valuable.

Applying `gramtools`’ genome graph model on microbial datasets, we obtain three main results. First, in *P. falciparum* genes with high diversity, `gramtools` geno-typing with genome graphs outperforms reference-genome-based callers, and these callers can use `gramtools`’ inferred personalised reference genome to access further variation. Second, `gramtools` provides superior genotyping accuracy compared to genome graph tools `vg` and `GraphTyper2` when jointly genotyping large deletions and overlapping small variants in *M. tuberculosis*. Note that for `GraphTyper2`, this use case is not directly supported, as it is designed to genotype structural variants only -we include the results anyway as this outperformed `vg`. We also note that during the finalisation of this paper, a new caller based on `vg` (named `Giraffe` [8]) was released, which we have not tested here. Third, we show how locally defined alternate references allow accessing small variants on top of diverged forms of a dimorphic gene in *P. falciparum*.

These results highlight three central concepts for genome graph based analyses: compatibility, consistency and interpretability.

First, while genome graphs extend beyond a single linear reference, maintaining compatibility with linear references is essential. `gramtools` outputs variation in terms of the standard reference genome in a VCF file and also produces a personalised reference genome, allowing reference-based callers to discover previously inaccessible variation. Many genomic analyses rely on a linear reference, which provides a simple coordinate system for referring to genomic annotations and comparing individuals. Recently the rGFA format for describing genome graphs was proposed [34]; starting from a central linear reference it assigns stable names and offset coordinates to alternate references. rGFA is a valuable and complementary idea to the jVCF described here: it assigns coordinates and references in constructed genome graphs, while our jVCF describes sites and called variation in genotyped genome graphs. Haplogroups output in the jVCF correspond to dynamically-defined alternate references; where useful these can be replaced with stable rGFA references defined at graph construction time. Like the NDAGs used by `gramtools`, rGFA works on globally linear graphs in order to maintain clear homology relationships.

Second, genome graphs offer the opportunity to genotype cohorts of samples consistently. By representing all variation found in a set of samples, they can be used to produce a full sample by site matrix. `gramtools` achieves this by detecting all variant sites in the graph and outputting them, along with their relationships, in a jVCF call format. Previous work has explored graph decomposition into a fixed set of variant sites [35] and is available in `vg` with the *deconstruct* command. However `vg` genotyping currently does not output all such sites nor define and output alternate references. `gramtools` provides the first working implementation of consistent graph to variant site mapping.

An important determinant of compatibility and consistency is the graph construction process. In `gramtools` we use our tool `make prg` [14]. From a multiple sequence alignment, `make prg` collapses common sequence between samples, clusters the remaining sequence into subgroups, and repeats the process recursively. This algorithm provides two main advantages. First, it limits recombination to similar input haplotypes, which reduces combinatorial explosions in variant dense regions, a source of computational bottlenecks and graph ambiguity [36]. Second, it naturally creates a hierarchy between sites as they are gradually defined on different sequence backgrounds. This captures incompatibility between sites (as in SNPs under a large deletion) as well as the process of divergent sequence evolution.

Finally, while single references and VCF provide good interpretability, we show how analysing two diverged forms of a dimorphic surface antigen in *P. falciparum* (DBLMSP2) benefits from locally defined alternate references. In contrast to existing sequences such as the alternate MHC loci in the human reference genome [37], here these are tied together in a graph-based framework. Outputting variation on different sequence backgrounds can provide finer resolution than with a single reference and will enable studying the functional impact and population genetics of nested variants.

## Conclusions

We provide a framework for identifying and genotyping multiscale variation in genome graphs and show its successful implementation in `gramtools`. We find good genotyping performance compared to state-of-the-art genome graph tools `GraphtTyper2` and `vg`, and additionally provide an analysis of allelic dimorphism using multiple references which to our knowledge can only be performed by `gramtools`.

Multiscale variation analysis goes hand in hand with the gradual extension of reference genomes beyond their linear coordinates. Accessing this complex variation requires careful genome graph construction and stable names and coordinates for referring to alternate references. It also calls for new developments in variant call output formats, a proposal of which we implement and use in `gramtools`.

## Methods

### Graph definitions

Here we formally define a variant site and the type of graph that gramtools can support. Let *G* = (*V, E*) be a directed acyclic graph (DAG) with a unique minimal and unique maximal element, ie *G* has a unique source and unique sink. Each node *v* has a number of ingoing edges *deg*^−^(*v*) and a number of outgoing edged *deg*^+^(*v*). Define a node *v* to be opening if *deg*^+^(*v*) *>* 1 and closing if *deg*^−^(*v*) *>* 1. Note that a node can be both opening and closing.

Given any opening node *v*, let *S* be the set of nodes that are in every path from *v* to the sink, excluding *v* itself. Then *S* is non-empty (because the sink is in *S*) and is totally ordered, and therefore has a minimal element, which we denote *c*(*v*). Informally, it can be thought of as the first node that “closes” all paths from *v*. Similarly, given a closing node u, we define *o*(*u*) to be *c*(*u*) applied to the transpose of *G*. Informally, *o*(*u*) is the node that “opened” *u*.

**Definition** *Variant site. A variant site in G is defined as the subgraph induced from* {*u, c*(*u*)} *or from* {*o*(*v*), *v*}, *where u is any opening node and v is any closing node*.

`gramtools` supports a DAG with a unique source, a unique sink, and satisfying the following property:

**Definition** *Balanced brackets property. There exists a topological ordering of all nodes v*_0_, *v*_1_, …, *v*_*n*_ *such that adding brackets to this ordered list of nodes according to the following rules results in balanced opening and closing brackets:*

1. *For each opening node u, add* (_*u*_ *after u, and add*)_*u*_ *before c*(*u*)
2. *For each closing node v, add* (_*o*(*v*)_ *after o*(*v*) *and add*)_*o*(*v*)_ *before v, unless these brackets were already added by case 1*

Each matching pair of brackets in this representation corresponds to one variant site. See Supplementary Fig. 2 for an illustration.

The above text fully describes the topology of sequence graphs supported by gramtools. However, to be able to index with the vBWT, `gramtools` also applies a simple transformation. For each matching pair of brackets *p* = {(_*u*_,)_*v*_} where *u* and *v* are opening and closing nodes of a variant site, and starting from the innermost bracket pairs:

- Add a node *s*_1_ to the graph, with no sequence, making a directed edge from *u* to *s*_1_ and moving *u*’s outgoing edges to *s*_1_
- Add a node *s*_2_ to the graph, with no sequence, making a directed edge from *s*_2_ to *v* and moving *v*’s incoming edges to *s*_2_

Now, each variant site in the graph has a pair of nodes {*s*_1_, *s*_2_} as opening and closing nodes, and each variant site satisfies the following property:

**Definition** *Disjoint subgraph property. Given a variant site with opening and closing nodes* {*s*_1_, *s*_2_}, *partition all walks from s*_1_ *into n sets S*_*i*_, *i* ∈ {1 … *n*} *of walks based on which of the n possible next nodes they move to after s*_1_. *If we let N* (*S*_*i*_) *denote the set of all nodes touched by all walks in set S*_*i*_, *then N* (*S*_*i*_) ⋂*N* (*S*_*j*_) = {*s*_1_, *s*_2_} *for all* {*i, j*} ∈ {1, *n*}. *In other words, the walks that spread out from s*_1_ *traverse completely disjoint subgraphs until they meet at s*_2_.

See Supplementary Fig. 3 for an illustration. We call graphs supported by `gramtools` nested DAGs (NDAGs), as variant sites can only overlap if one is fully contained (nested) within another.

### vBWT data structure in gramtools

The vBWT data structure marks variant sites with numeric identifiers so that alleles get sorted and queried together in the suffix array (Fig. 9 panel a). This representation induces branching at each site entry and exit such that mapping has worst-case exponential run-time. To speed mapping, we seed reads from an index storing the mapped intervals of all sequences of a given size *k*. Linear-time exact match indexes on genome graphs exist (e.g. GCSA [38]) but require a prefix-sorting step that is worst-case exponential.

**Figure 9.**
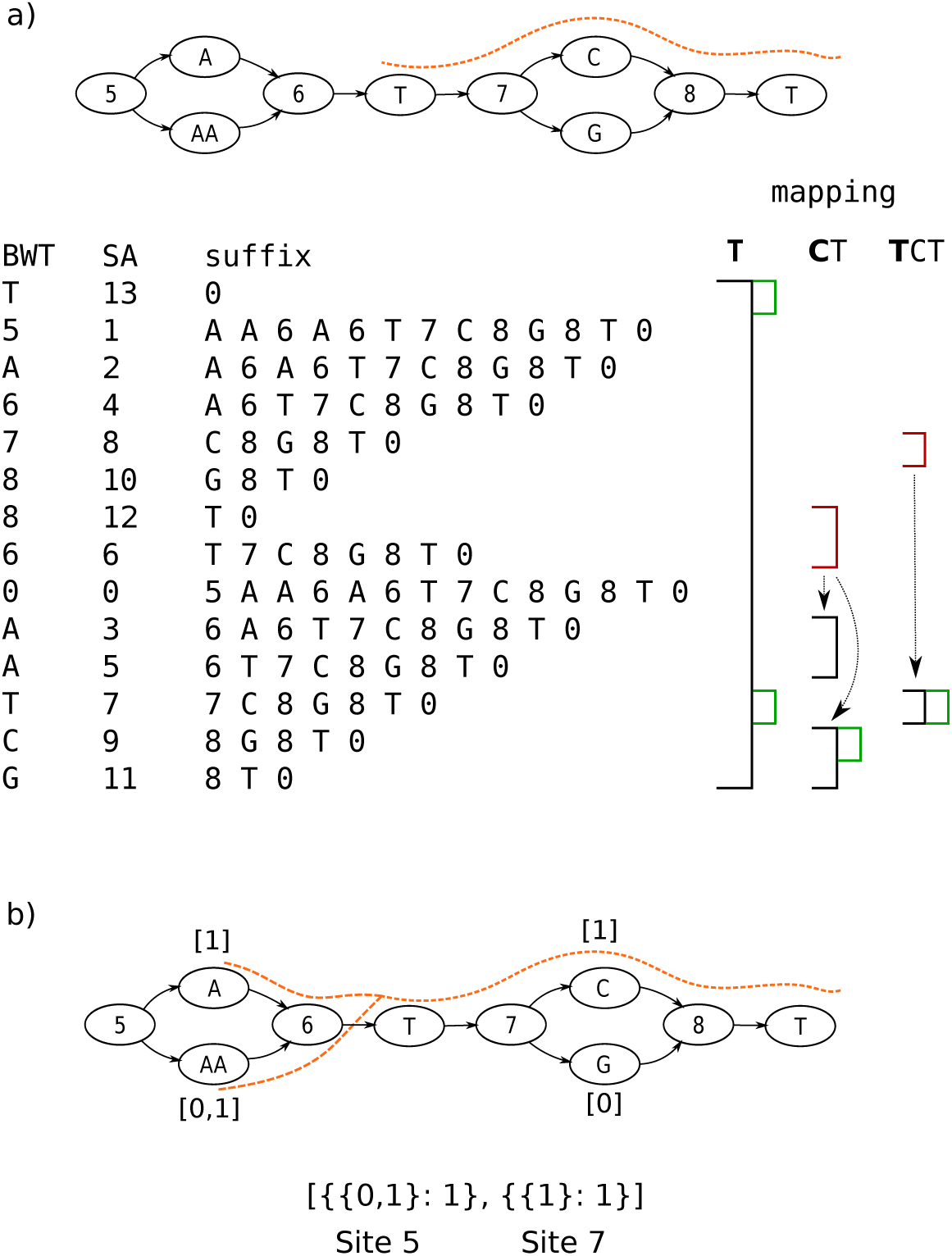
gramtools mapping and coverage recording. a) variant-aware Burrows Wheeler Transform (vBWT). Each row of the text matrix encodes one position in a linear representation of the graph. BWT: stores the character in the previous position; SA: suffix array, stores the position in the text; suffix: stores the text from SA position to the end. Two markers are used for every variant site in the genome graph: odd markers mark site entry and even markers allow alleles to sort and be queried together. Black intervals mark regular BWT backward searching, with each match to the currently mapped base shown in green. Arrows from red intervals mark vBWT-specific jumps in and out of sites, making the search branch. The read being mapped is shown in dashed orange. b) Square brackets under allele nodes show per-base coverage storage. Another array shown below stores allele-level coverage at each site. Mapped reads increment equivalence class counts representing compatibility: in this example, the read is compatible with both alleles 0 and 1 at site 5 and only with allele 1 at site 7. Both kinds of coverage are used in genotyping.

vBWT’s numeric identifiers are also used for recording mapped read coverage along variant sites (Fig.9, Panel b). Coverage recording handles two types of uncertainty: horizontal, where sequence is repeated across the genome, and vertical, where sequence is repeated in alleles of a site. To handle horizontal uncertainty we randomly select one read mapping instance, as is typically done in standard aligners [30]. To handle vertical uncertainty we store allele-level equivalence class counts which are counts of reads compatible with groups of alleles, an idea introduced in `kallisto` [15]. This allows allelic uncertainty to be accounted for during genotyping. Per-base coverage is also stored on the graph (Fig.9) and used during genotyping.

### Genotyping model

The genotyping model in `gramtools` supports haploid and diploid genotyping. It assigns a likelihood to each candidate allele (or pair of alleles for diploid) computed from base-level and allele-level read coverage.

`gramtools` stores coverage in equivalence classes, following ideas from [15]. Let *A* be the set of alleles at a variant site. We partition the set of all reads overlapping *A* into subsets, i.e. equivalence classes, where all reads belonging to one subset map perfectly to the same subset *X* of *A* (e.g. reads that map uniquely to allele 1, or reads that map equally well to alleles 1 and 2; see Fig. 9 Panel b). For each equivalence class, we store a count *c*_*X*_ ∈ ℕ of reads compatible with the alleles in set *X*, and for each mapped read we increment its *c*_*X*_ at each overlapped site. If a read has multiple (horizontal) mapping instances we select one at random, and the counts *c*_*X*_ are incremented as above. When a read’s count *c*_*X*_ is incremented, for each allele *a* ∈ *X* the count of each base the read mapped to is also incremented. Base-level counts are written *c*(*a*_*i*_), where *a*_*i*_ is the *i*th base of allele *a*.

Coverage compatible with allele *a* of length *l*_*a*_ is defined as 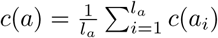 ,and incompatible coverage as *i*(*a*) = ∑_X ⊂A:*a* ∉X_ *c*XIn this way for each candidate allele we capture a per-base correct coverage generation process as well as an incorrect coverage generation process on incompatible alleles.

We model the expected per-base read coverage in a site using an estimate of the mean *λ* and the variance *σ*^2^ of true coverage across all variant sites. For each site true coverage is estimated as the average per-base coverage of the allele with the most coverage. If *σ*^2^ ⩽ *λ*, we model observed coverage as coming from a Poisson distribution:

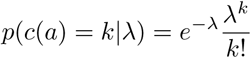

Else, we use the negative binomial (NB) distribution

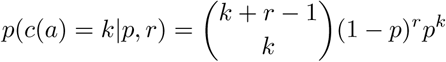

When using the NB distribution we need to estimate the parameters of the distribution. The number of failures *r* before *k* successes is estimated from the expected variance of NB as 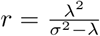 and the probability of success *p* is estimated from the expected mean of NB as 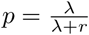

We model incorrect coverage *i*(*a*) as coming from sequencing errors with rate *ϵ*: *p*(*i*(*a*) = *k*|*ϵ*) = *ϵ*^*k*^. *ϵ* is estimated as the mean base quality score in the first 10,000 processed reads.

We also use per-base coverage to penalise gaps in coverage. Given a function *g*(*a*) returning the number of zero-coverage positions in allele *a*, the probability *p*(*g*(*a*) = *k*) of seeing *k* gaps if *a* is the true allele is *p*(*c*(*a*) = 0)^*k*^. In practice we use 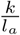 as the exponent so as not to penalise long alleles.

These three terms combined give the likelihood of allele *a*

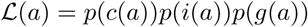

The allele *a*^′^ that gets called is the maximum-like9lihood allele and we define the genotype confidence of the call as

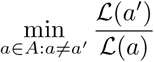

which is the likelihood ratio of the called allele and the next most likely allele.

This holds for haploid genotyping. For higher ploidy, the likelihood function generalises to a set of alleles *S* as

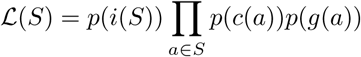

where *i*(*S*) = Σ_*X*⊂*A*:*X*∩*S*=∅_ *c*_*X*_.

For diploid genotyping, *S* = {*a*_1_, *a*_2_} and *p*(*c*(*a*)) is parameterised by 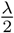 because we expect half the total site coverage on each of two called alleles.

### Nested genotyping

The `gramtools` genotying model is applied recursively from child sites to their parent sites. Calls in child sites restrict the set of alleles considered in the parent so that the number of choices is reduced: for each outgoing path from a site, 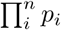 paths are considered, where *p*_*i*_ is the number of distinct alleles called at site *i* (e.g. in diploids 0, 1 or 2) and *n* the number of child sites encountered. Some extra paths are also retained when genotype confidence for a child site is low, in order to propagate uncertainty to parent calls. If there are more than 10,000 possible alleles, only the 10,000 most likely alleles are considered. This does not require enumerating all possible alleles as the most likely alleles in child sites have already been computed.

### Genome graph construction using make prg

`gramtools` can construct genome graphs without nested variation from a reference genome and a VCF of variants. Overlapping records in the VCF file are merged by enumerating all possible combinations up to a specifiable limit. This method rapidly fails in variant-dense regions or for large cohorts of samples and does not allow nested variation. To build nested graphs we apply an algorithm called recursive collapse and cluster (RCC) starting from a multiple-sequence alignment. RCC was first introduced in the context of bacterial pan-genomic tool `pandora` [14].

RCC identifies invariant regions of a given minimum size and collapses them together. The remaining regions are clustered based on their *k* -mer content. This procedure is repeated recursively on each cluster, until either a maximum nesting level is reached or the sequences are too small (in which case they are directly enumerated as alternative alleles). In this way variants appear in subsets of samples with similar sequence backgrounds and only share sequence if one occurs inside the other. The output graph is a nested, directed acyclic graph (NDAG), which is a requirement for `gramtools`.

Two command-line parameters affect what graph gets produced. First, *max nesting* is the maximum number of collapse and cluster recursions to perform, which gives the maximum number of nesting levels in the graph. Second, *min match length* is the number of invariant bases between samples for them to be collapsed in a single node. Sequence collapse is what allows paths coming before and after to cross; a larger value thus reduces recombination between the input haplotypes. This provides a way to control combinatorial path explosions in the graph.

RCC is implemented in and available at https://github.com/iqbal-lab-org/make_prg.

### *P. falciparum* surface antigen graphs and genotyping validation

#### Graph construction

We started from VCF files produced by running `Cortex`, a *de novo* assembly based variant caller [5], on read sets of 2,498 samples from the Pf3k project [19]. `Cortex` has a very low false positive call rate and can call the divergent forms of *P. falciparum* surface antigens [21]. The `Cortex` independent workflow was run, using the bubble caller with *k=31*. The VCF files are publicly released on zenodo (see availability) and `Cortex` is publicly available at https://github.com/iqbal-lab/cortex.

For each surface antigen gene (DBLMSP, DBLMSP2, EBA175 and AMA1), we generated sequences for each sample by applying `Cortex` variants at the gene coordinates, plus 5000bp on each side, to the *P. falciparum* 3D7 reference genome. We generated multiple-sequence alignments of each gene using `mafft` [39] and passed them as input to our construction tool `make prg`. For the simulation experiment, two graphs were built for DBLMSP and DBLMSP2 with maximum nesting levels of 1 and 5. The graphs without nesting have 451 and 413 variant sites for DBLMSP and DBLMSP2, and the graphs with nesting have 558 and 500 variant sites respectively.

#### Path and read simulation

From each non-nested graph, 10 paths were simulated and threaded through the nested graph using `gramtools`’ `simulate` command. This results in jVCF files recording the true genotypes for each path in each graph. Illumina HiSeq25 75-bp reads (0.023% per-base error rate) were simulated from each unique path using ART [40] at 40-fold coverage, genotyped using `gramtools` and calls evaluated by comparing the genotyped and truth jVCF files.

The nested graph contains more paths than the non-nested graphs due to allowing greater recombination between variant sites. We therefore simulated paths from the non-nested graph to ensure each path exists in both graphs.

#### Comparison with reference-based callers

The nested graphs of the four surface antigens DBLMSP, DBLMSP2, EBA175 and AMA1 were combined in positional order with the rest of the reference genome, using a custom script (see availability).

For each of the 14 validation samples, we ran `SAMtools` and `Cortex` using the *P. falciparum* 3D7 reference genome and `gramtools` using the surface antigen genome graph. For each genotyped sample, `gramtools` infers a haploid personalised genome (PR) as the whole-genome path taking the called allele at each variant site. `SAMtools` and `Cortex` were then run once more using the personalised reference instead of 3D7.

When mapping the gene sequence with variants applied to the truth assemblies, we measure performance as the edit distance reported by `bowtie2` (version 2.4.1) divided by gene length.

#### Determining the closest input panel sequence

To determine the closest panel sequence to a truth assembly, for each gene we mapped all 2,498 sequences used for building the gene’s graph to the truth assembly using `bowtie2`. The closest sequence was the one with the lowest ‘NM’ (edit distance) tag, provided it mapped with a MAPQ *>* 40 (i.e. the probability the mapping was not unique was *<* 10^−4^ according to `bowtie2`).

### *M. tuberculosis* SNP and large deletion analysis

#### Long-read assembly of the 17 evaluated samples

Each sample was initially assembled using `Unicycler` [41] and `Canu` [42], followed by `Circlator` [43] using the corrected reads output by `Canu`. `Unicycler` version 0.4.8 was used with the option ‘`-mode bold`’, Illumina reads given using the options ‘`--short1` and ‘`--short2`’, and the PacBio subreads using the ‘`--long`’ option. `Canu` version 2.0 was used with the option ‘`genomeSize=4.4m`’ and the PacBio reads provided with the option ‘`-pacbio-raw`’. The only exception was sample N1177, which was initially assembled using `Flye` [44] version 2.8-b1674 with the PacBio subreads input with the option ‘`--pacbio-raw`’.

The initial assembly for each sample was chosen for further manual polishing based on inspection of mapped reads and comparison with the H37Rv reference genome. The `Unicycler` assembly was used for samples N0004, N0091, N0155, N0157, N1283, N0072, N1202, N0153. The Canu assembly was used for samples N0031, N1176, N0052, N0136, N0146, N1216, N1272, N0054. Redundant and/or contamination contigs were removed from samples N0072, N1202, N0052, N0136, N0146, N1216, N1272. Manual fixes were applied to samples N0054 and N0153 by breaking contigs at errors, with the aid of the Artemis Comparison Tool (`ACT`) [47], and re-merging using `Circlator` using the default settings. Next, `Pilon` [45] (version 1.23) was run iteratively on each assembly using the Illumina reads as input, mapped with `BWA-MEM` [46] (version 0.7.17-r1188, default settings) until no more corrections were made, up to a maximum of 10 iterations. Finally, the ‘fixstart’ function of `Circlator` was used to ensure that each assembly began with the *dnaA* gene, for consistency with the H37Rv reference genome. The result was all 17 samples assembled into a single, circularized contig.

#### Variant discovery

We first obtained variant calls from the read files of the 17 evaluated samples and an additional 1,000 samples available in the ENA (see availability). We ran `Cortex` to obtain the calls, using our lab’s wrapper `clockwork` version 0.8.3, publicly available at https://github.com/iqbal-lab-org/clockwork. `clockwork` runs `Cortex`’s in-dependent workflow using the bubble caller with *k=31*. The VCF files are publicly released on zenodo (see availability).

`Cortex` identified a total of 73 deletions in the 17 evaluated samples, between 100 and 13,000 bases in length and falling in 45 distinct genomic regions. To validate the calls, we mapped their corresponding long-read assemblies to the *M. tuberculosis* H37Rv reference genome with `minimap2`, which validated 68. The remaining 5 were manually confirmed using `ACT`: for each sample we mapped the short reads to the reference genome and to the assembly using `bowtie2`, and mapped the assembly to the reference using `nucmer` [48]. In `ACT` we view all three together and validate a deletion when it appears in the assembly-reference mapping at the expected coordinates and when read pileups confirm the event. An example is shown in Supplementary Fig. 11.

Having validated all the deletions, we extracted all `Cortex` calls occurring under the 45 deletion regions in the 1,017 samples, giving us a joint set of large deletions and overlapping SNPs and indels.

#### gramtools genome graph construction

We built one genome graph for each of the 45 regions identified as containing large deletions in our 17 evaluation samples. As for the *P. falciparum* surface antigen graphs, for each region we applied `Cortex` calls to the *M. tuberculosis* H37Rv reference genome, generated multiple sequence alignments with `mafft`, passed them as input to our construction tool `make prg` and combined them with the rest of the genome.

#### vgand GraphtTyper2 genome graph construction

We set on building a `vg` genome graph from the same multiple sequence alignments (MSA) used by `gramtools` to maximise comparability. Using `vg` version 1.26.0, we built each of the 45 regions from MSA using `vg construct` and combined them with the invariant parts of the *M. tuberculosis* H37Rv reference genome using `vg concat`.

Indexing this graph, a prerequisite to read mapping and variant calling, used *>*10 Terabytes of temporary disk space before we stopped it. We deemed *>*500 Gigabytes prohibitive and set that as a limit. We ran `vg prune` to remove densely clustered variation from the graph and, after exceeding 1 Terabyte of disk indexing the pruned graph using default parameters, successfully indexed the pruned graph with parameters `-k10 -X3`.

We then ran `vg call` for each of our 1,017 samples against the MSA graph. However, after successful mapping to this indexed graph, `vg call` failed with a segmentation fault.

We therefore built a graph from a VCF file instead. We ran `vg deconstruct –p -e` to obtain a VCF file describing the variants identified by `vg` in the `vg` MSA-constructed graph, and manually validated the variation using one sample when compared to `gramtools`. However running `vg construct` with this VCF also failed with a segmentation fault.

We therefore used `vg` graph construction and genotyping from a merged VCF of all variants in the 45 regions which we produced using `bcftools`. This ran successfully after graph pruning to stay under our disk limit.

This VCF file was also used as input to `GraphtTyper2` via `graphtyper` version 2.5.1, running its `genotype sv` subcommand. `GraphtTyper2` only accepts VCF files as input and not MSAs.

#### Mapping evaluated regions to truth assemblies

We evaluated a total of 3,060 sequences by mapping them to truth assemblies: 17 samples x 45 regions x 4 tools (`gramtools`, `vg`, `GraphtTyper2` and the reference genome sequence). Using `bowtie2` 10.4% all sequences failed to be fully aligned due to excessive divergence between the called sequence and the truth.

To recover more alignments, we used `minimap2` which is designed to align more highly diverged sequences (such as ONT long reads) [29]. For each evaluated sequence, we took the alignment with the greatest number of matches to the assembly and extracted assembly sequence of the same length from the first aligned position (including soft-or hard-clipped). We obtained the edit distance between the two sequences from Needleman-Wunsch alignment using `edlib` [49]. Using this approach reduced the proportion of unaligned sequences to 1.1%.

To ensure evaluated alignments are unambiguous, we filter them by MAPQ ≥30 so that the probability they are non-unique is ≤10^−3^ as estimated by `minimap2`. This removed 0.62% of the evaluated sequences. For each tool, 13 of 765 sequences were not mapped or had insufficiently high mapping quality. The number of unmapped and low MAPQ sequences for each tool are shown in Supplementary Fig. 12.

We required the VCF records output by each tool to have a FILTER status set to “PASS”. This changed results only marginally, giving the same number of unmapped and low MAPQ sequences and a decreased mean edit distance by 0.11% for `GraphtTyper2`, 0.08% for `gramtools` and no differences for `vg`.

### *P. falciparum* dimorphic variation analysis

We obtained read files for 706 samples from the Pf3k project [19]. The samples come from Ghana, Cambodia and Laos and are publicly available on the ENA (see availability). Each sample was genotyped on the previously-built *P. falciparum* graph of four surface antigen genes (DBLMSP, DBLMSP2, EBA175 and AMA1) (see Graph construction).

Using `gramtools`, we combined the per-sample jVCF output files into one multi-sample jVCF file, which we analysed using custom scripts (see availability).

## Authors’ contributions

BL developed `gramtools`, performed the analyses, drafted the manuscript and wrote the repository for the study (see availability). MH produced the *M. tuberculosis* assemblies and Cortex VCF files and helped using `ACT`. ZI designed the study, produced the *P. falciparum* Cortex VCF files. MH and ZI defined the graph constraints and edited the manuscript. MH and ZI designed, and BL modified, the genotyping model. All authors reviewed and approved the manuscript.

## Acknowledgements

The authors thank Rachel Colquhoun for the algorithms and first development of make prg, Sorina Maciuca for the vBWT data structure and algorithms behind gramtools, and Robyn Ffrancon for software engineering in gramtools.

## Availability of data and materials

gramtools is open-source and available at https://github.com/iqbal-lab-org/gramtools. We provide an open repository for reproducing all results in this study available at https://github.com/iqbal-lab-org/paper_gramtools_nesting. Each part of the analysis can be rerun using the Snakemake workflow engine [50].

Data is available via the repository in two forms. Accessions for data deposited at the ENA are stored as tables. All other data, as well as a software container, have been released for this study on zenodo at https://doi.org/10.5281/zenodo.4147302. All versions/commits of the software used in this study are frozen in the container and can be found at

https://github.com/iqbal-lab-org/paper_gramtools_nesting/blob/master/container/singu_def.def. The repository README provides all the instructions to obtain the data and to re-run each part of the analysis.

## Ethics approval and consent to participate

Not applicable.

## Author details

EMBL-EBI, Hinxton, UK.

## References

1. Brandt, D.Y.C., Aguiar, V.R.C., Bitarello, B.D., Nunes, K., Goudet, J., Meyer, D.: Mapping bias overestimates reference allele frequencies at the hla genes in the 1000 genomes project phase i data. G3 (Bethesda, Md.) 5(5), 931–941 (2015). doi:10.1534/g3.114.015784

2. Schneeberger, K., Hagmann, J., Ossowski, S., Warthmann, N., Gesing, S., Kohlbacher, O., Weigel, D.: Simultaneous alignment of short reads against multiple genomes. Genome Biology 10(9), 98 (2009). doi:10.1186/gb-2009-10-9-r98

3. Dilthey, A., Cox, C., Iqbal, Z., Nelson, M.R., McVean, G.: Improved genome inference in the MHC using a population reference graph. Nature Genetics 47(6), 682–688 (2015). doi:10.1038/ng.3257

4. Garrison, E., Sirén, J., Novak, A.M., Hickey, G., Eizenga, J.M., Dawson, E.T., Jones, W., Garg, S., Markello, C., Lin, M.F., Paten, B., Durbin, R.: Variation graph toolkit improves read mapping by representing genetic variation in the reference. Nature Biotechnology 36(9), 875–879 (2018). doi:10.1038/nbt.4227

5. Iqbal, Z., Caccamo, M., Turner, I., Flicek, P., McVean, G.: De novo assembly and genotyping of variants using colored de Bruijn graphs. Nature Genetics 44(2), 226–232 (2012). doi:10.1038/ng.1028

6. Eggertsson, H.P., Jonsson, H., Kristmundsdottir, S., Hjartarson, E., Kehr, B., Masson, G., Zink, F., Hjorleifsson, K.E., Jonasdottir, A., Jonasdottir, A., Jonsdottir, I., Gudbjartsson, D.F., Melsted, P., Stefansson, K., Halldorsson, B.V.: Graphtyper enables population-scale genotyping using pangenome graphs. Nature Genetics 49(11), 1654–1660 (2017). doi:10.1038/ng.3964

7. Sibbesen, J.A., Maretty, L., Krogh, A.: Accurate genotyping across variant classes and lengths using variant graphs. Nature Genetics 50(7), 1054 (2018). doi:10.1038/s41588-018-0145-5

8. Sirén, J., Monlong, J., Chang, X., Novak, A.M., Eizenga, J.M., Markello, C., Sibbesen, J., Hickey, G., Chang, P.-C., Carroll, A., Haussler, D., Garrison, E., Paten, B.: Genotyping common, large structural variations in 5,202 genomes using pangenomes, the giraffe mapper, and the vg toolkit. bioRxiv (2020). doi:10.1101/2020.12.04.412486. https://www.biorxiv.org/content/early/2020/12/06/2020.12.04.412486.full.pdf

9. Eggertsson, H.P., Kristmundsdottir, S., Beyter, D., Jonsson, H., Skuladottir, A., Hardarson, M.T., Gudbjartsson, D.F., Stefansson, K., Halldorsson, B.V., Melsted, P.: GraphTyper2 enables population-scale genotyping of structural variation using pangenome graphs. Nature Communications 10(1), 5402 (2019). doi:10.1038/s41467-019-13341-9. Number: 1 Publisher: Nature Publishing Group

10. Danecek, P., Auton, A., Abecasis, G., Albers, C.A., Banks, E., DePristo, M.A., Handsaker, R.E., Lunter, G., Marth, G.T., Sherry, S.T., McVean, G., Durbin, R., Group, .G.P.A.: The variant call format and VCFtools. Bioinformatics 27(15), 2156–2158 (2011). doi:10.1093/bioinformatics/btr330. Publisher: Oxford Academic

11. Maciuca, S., Elias, C.d.O., McVean, G., Iqbal, Z.: A natural encoding of genetic variation in a Burrows-Wheeler Transform to enable mapping and genome inference. In: Springer (ed.) Proceedings of the 16th International Workshop on Algorithms in Bioinformatics, Volume 9838 of Lecture Notes in Computer Science, pp. 222–233 (2016)

12. Valenzuela, D., Norri, T., Välimäki, N., Pitkänen, E., Mäkinen, V.: Towards pan-genome read alignment to improve variation calling. BMC Genomics 19(2), 87 (2018). doi:10.1186/s12864-018-4465-8

13. Ferragina, P., Manzini, G.: Opportunistic data structures with applications. In: Proceedings 41st Annual Symposium on Foundations of Computer Science, pp. 390–398. IEEE Comput. Soc, Redondo Beach, CA, USA (2000). doi:10.1109/SFCS.2000.892127. http://ieeexplore.ieee.org/document/892127/

14. Colquhoun, R.M., Hall, M.B., Lima, L., Roberts, L.W., Malone, K.M., Hunt, M., Letcher, B., Hawkey, J., George, S., Pankhurst, L., Iqbal, Z.: Nucleotide-resolution bacterial pan-genomics with reference graphs. bioRxiv, 2020–1112380378 (2020). doi:10.1101/2020.11.12.380378. Publisher: Cold Spring Harbor Laboratory Section: New Results

15. Bray, N.L., Pimentel, H., Melsted, P., Pachter, L.: Near-optimal probabilistic RNA-seq quantification. Nature Biotechnology 34(5), 525–527 (2016). doi:10.1038/nbt.3519

16. Ochola, L.I., Tetteh, K.K.A., Stewart, L.B., Riitho, V., Marsh, K., Conway, D.J.: Allele Frequency–Based and Polymorphism-Versus-Divergence Indices of Balancing Selection in a New Filtered Set of Polymorphic Genes in Plasmodium falciparum. Molecular Biology and Evolution 27(10), 2344–2351 (2010). doi:10.1093/molbev/msq119. Publisher: Oxford Academic

17. Amambua-Ngwa, A., Tetteh, K.K.A., Manske, M., Gomez-Escobar, N., Stewart, L.B., Deerhake, M.E., Cheeseman, I.H., Newbold, C.I., Holder, A.A., Knuepfer, E., Janha, O., Jallow, M., Campino, S., MacInnis, B., Kwiatkowski, D.P., Conway, D.J.: Population Genomic Scan for Candidate Signatures of Balancing Selection to Guide Antigen Characterization in Malaria Parasites. PLOS Genetics 8(11), 1002992 (2012). doi:10.1371/journal.pgen.1002992

18. Maciuca, S.: Analysis of complex genetic variation using population reference graphs. PhD thesis, University of Oxford (2017)

19. The Pf3K Project (2015): Pilot Data Release 3. http://www.malariagen.net/data/pf3k-3

20. Li, H., Handsaker, B., Wysoker, A., Fennell, T., Ruan, J., Homer, N., Marth, G., Abecasis, G., Durbin, R., Subgroup, .G.P.D.P.: The Sequence Alignment/Map format and SAMtools. Bioinformatics 25(16), 2078–2079 (2009). doi:10.1093/bioinformatics/btp352. https://academic.oup.com/bioinformatics/article-pdf/25/16/2078/531810/btp352.pdf

21. Miles, A., Iqbal, Z., Vauterin, P., Pearson, R., Campino, S., Theron, M., Gould, K., Mead, D., Drury, E., O’Brien, J., Rubio, V.R., MacInnis, B., Mwangi, J., Samarakoon, U., Ranford-Cartwright, L., Ferdig, M., Hayton, K., Su, X.-z., Wellems, T., Rayner, J., McVean, G., Kwiatkowski, D.: Indels, structural variation, and recombination drive genomic diversity in Plasmodium falciparum. Genome Research 26(9), 1288–1299 (2016). doi:10.1101/gr.203711.115

22. Richards, J.S., Beeson, J.G.: The future for blood-stage vaccines against malaria. Immunology & Cell Biology 87(5), 377–390 (2009). doi:10.1038/icb.2009.27. https://onlinelibrary.wiley.com/doi/pdf/10.1038/icb.2009.27

23. Barry, A.E., Arnott, A.: Strategies for designing and monitoring malaria vaccines targeting diverse antigens 5. doi:10.3389/fimmu.2014.00359

24. Otto, T.D., Böhme, U., Sanders, M., Reid, A., Bruske, E.I., Duffy, C.W., Bull, P.C., Pearson, R.D., Abdi, A., Dimonte, S., Stewart, L.B., Campino, S., Kekre, M., Hamilton, W.L., Claessens, A., Volkman, S.K., Ndiaye, D., Amambua-Ngwa, A., Diakite, M., Fairhurst, R.M., Conway, D.J., Franck, M., Newbold, C.I., Berriman, M.: Long read assemblies of geographically dispersed Plasmodium falciparum isolates reveal highly structured subtelomeres. Wellcome Open Research 3 (2018). doi:10.12688/wellcomeopenres.14571.1

25. Gardner, M.J., Hall, N., Fung, E., White, O., Berriman, M., Hyman, R.W., Carlton, J.M., Pain, A., Nelson, K.E., Bowman, S., Paulsen, I.T., James, K., Eisen, J.A., Rutherford, K., Salzberg, S.L., Craig, A., Kyes, S., Chan, M.-S., Nene, V., Shallom, S.J., Suh, B., Peterson, J., Angiuoli, S., Pertea, M., Allen, J., Selengut, J., Haft, D., Mather, M.W., Vaidya, A.B., Martin, D.M.A., Fairlamb, A.H., Fraunholz, M.J., Roos, D.S., Ralph, S.A., McFadden, G.I., Cummings, L.M., Subramanian, G.M., Mungall, C., Venter, J.C., Carucci, D.J., Hoffman, S.L., Newbold, C., Davis, R.W., Fraser, C.M., Barrell, B.: Genome sequence of the human malaria parasite plasmodium falciparum 419(6906). doi:10.1038/nature01097

26. Borrell, S., Trauner, A., Brites, D., Rigouts, L., Loiseau, C., Coscolla, M., Niemann, S., De Jong, B., Yeboah-Manu, D., Kato-Maeda, M., Feldmann, J., Reinhard, M., Beisel, C., Gagneux, S.: Reference set of Mycobacterium tuberculosis clinical strains: A tool for research and product development. PLoS ONE 14(3) (2019). doi:10.1371/journal.pone.0214088

27. Chiner-Oms, A., Berney, M., Boinett, C., González-Candelas, F., Young, D.B., Gagneux, S., Jacobs, W.R., Parkhill, J., Cortes, T., Comas, I.: Genome-wide mutational biases fuel transcriptional diversity in the Mycobacterium tuberculosis complex. Nature Communications 10(1), 3994 (2019). doi:10.1038/s41467-019-11948-6

28. Cole, S.T., Brosch, R., Parkhill, J., Garnier, T., Churcher, C., Harris, D., Gordon, S.V., Eiglmeier, K., Gas, S., Barry, C.E., Tekaia, F., Badcock, K., Basham, D., Brown, D., Chillingworth, T., Connor, R., Davies, R., Devlin, K., Feltwell, T., Gentles, S., Hamlin, N., Holroyd, S., Hornsby, T., Jagels, K., Krogh, A., McLean, J., Moule, S., Murphy, L., Oliver, K., Osborne, J., Quail, M.A., Rajandream, M.-A., Rogers, J., Rutter, S., Seeger, K., Skelton, J., Squares, R., Squares, S., Sulston, J.E., Taylor, K., Whitehead, S., Barrell, B.G.: Deciphering the biology of mycobacterium tuberculosis from the complete genome sequence. Nature 393(6685), 537–544 (1998). doi:10.1038/31159

29. Li, H.: Minimap2: pairwise alignment for nucleotide sequences. Bioinformatics 34(18), 3094–3100 (2018). doi:10.1093/bioinformatics/bty191. Publisher: Oxford Academic

30. Langmead, B., Salzberg, S.L.: Fast gapped-read alignment with Bowtie 2. Nature Methods 9(4), 357–359 (2012). doi:10.1038/nmeth.1923

31. Büchler, T., Ohlebusch, E.: An improved encoding of genetic variation in a Burrows–Wheeler transform. Bioinformatics 36(5), 1413–1419 (2020). doi:10.1093/bioinformatics/btz782. Publisher: Oxford Academic

32. Crosnier, C., Iqbal, Z., Knuepfer, E., Maciuca, S., Perrin, A.J., Kamuyu, G., Goulding, D., Bustamante, L.Y., Miles, A., Moore, S.C., Dougan, G., Holder, A.A., Kwiatkowski, D.P., Rayner, J.C., Pleass, R.J., Wright, G.J.: Binding of Plasmodium falciparum Merozoite Surface Proteins DBLMSP and DBLMSP2 to Human Immunoglobulin M Is Conserved among Broadly Diverged Sequence Variants. Journal of Biological Chemistry 291(27), 14285–14299 (2016). doi:10.1074/jbc.M116.722074

33. Chen, X., Schulz-Trieglaff, O., Shaw, R., Barnes, B., Schlesinger, F., Källberg, M., Cox, A.J., Kruglyak, S., Saunders, C.T.: Manta: rapid detection of structural variants and indels for germline and cancer sequencing applications. Bioinformatics (Oxford, England) 32(8), 1220–1222 (2016). doi:10.1093/bioinformatics/btv710

34. Li, H., Feng, X., Chu, C.: The design and construction of reference pangenome graphs with minigraph. Genome Biology 21(1), 265 (2020). doi:10.1186/s13059-020-02168-z

35. Paten, B., Eizenga, J.M., Rosen, Y.M., Novak, A.M., Garrison, E., Hickey, G.: Superbubbles, Ultrabubbles, and Cacti. Journal of Computational Biology 25(7), 649–663 (2018). doi:10.1089/cmb.2017.0251. Publisher: Mary Ann Liebert, Inc., publishers

36. Pritt, J., Chen, N.-C., Langmead, B.: FORGe: prioritizing variants for graph genomes. Genome Biology 19(1), 220 (2018). doi:10.1186/s13059-018-1595-x

37. Church, D.M., Schneider, V.A., Steinberg, K.M., Schatz, M.C., Quinlan, A.R., Chin, C.-S., Kitts, P.A., Aken, B., Marth, G.T., Hoffman, M.M., Herrero, J., Mendoza, M.L.Z., Durbin, R., Flicek, P.: Extending reference assembly models. Genome Biology 16(1), 13 (2015). doi:10.1186/s13059-015-0587-3

38. Siren, J., Välimäki, N., Mäkinen, V.: [GCSA]Indexing Graphs for Path Queries with Applications in Genome Research. IEEE/ACM Transactions on Computational Biology and Bioinformatics 11(2), 375–388 (2014). doi:10.1109/TCBB.2013.2297101

39. Katoh, K., Misawa, K., Kuma, K., Miyata, T.: MAFFT: a novel method for rapid multiple sequence alignment based on fast Fourier transform. Nucleic Acids Research 30(14), 3059–3066 (2002). doi:10.1093/nar/gkf436. https://academic.oup.com/nar/article-pdf/30/14/3059/9488148/gkf436.pdf

40. Huang, W., Li, L., Myers, J.R., Marth, G.T.: ART: a next-generation sequencing read simulator. Bioinformatics 28(4), 593–594 (2011). doi:10.1093/bioinformatics/btr708

41. Wick, R.R., Judd, L.M., Gorrie, C.L., Holt, K.E.: Unicycler: Resolving bacterial genome assemblies from short and long sequencing reads. PLOS Computational Biology 13(6), 1005595 (2017). doi:10.1371/journal.pcbi.1005595. Publisher: Public Library of Science

42. Koren, S., Walenz, B.P., Berlin, K., Miller, J.R., Bergman, N.H., Phillippy, A.M.: Canu: scalable and accurate long-read assembly via adaptive k-mer weighting and repeat separation. Genome Research 27(5), 722–736 (2017). doi:10.1101/gr.215087.116. Company: Cold Spring Harbor Laboratory Press Distributor: Cold Spring Harbor Laboratory Press Institution: Cold Spring Harbor Laboratory Press Label: Cold Spring Harbor Laboratory Press Publisher: Cold Spring Harbor Lab

43. Hunt, M., Silva, N.D., Otto, T.D., Parkhill, J., Keane, J.A., Harris, S.R.: Circlator: automated circularization of genome assemblies using long sequencing reads. Genome Biology 16(1), 294 (2015). doi:10.1186/s13059-015-0849-0

44. Kolmogorov, M., Yuan, J., Lin, Y., Pevzner, P.A.: Assembly of long, error-prone reads using repeat graphs. Nature Biotechnology, 1 (2019). doi:10.1038/s41587-019-0072-8

45. Walker, B.J., Abeel, T., Shea, T., Priest, M., Abouelliel, A., Sakthikumar, S., Cuomo, C.A., Zeng, Q., Wortman, J., Young, S.K., Earl, A.M.: Pilon: An Integrated Tool for Comprehensive Microbial Variant Detection and Genome Assembly Improvement. PLOS ONE 9(11), 112963 (2014). doi:10.1371/journal.pone.0112963. Publisher: Public Library of Science

46. Li, H.: Aligning sequence reads, clone sequences and assembly contigs with BWA-MEM (2013). 1303.3997

47. Carver, T.J., Rutherford, K.M., Berriman, M., Rajandream, M.-A., Barrell, B.G., Parkhill, J.: ACT: the Artemis comparison tool. Bioinformatics 21(16), 3422–3423 (2005). doi:10.1093/bioinformatics/bti553. https://academic.oup.com/bioinformatics/article-pdf/21/16/3422/573752/bti553.pdf

48. Kurtz, S., Phillippy, A., Delcher, A.L., Smoot, M., Shumway, M., Antonescu, C., Salzberg, S.L.: Versatile and open software for comparing large genomes. Genome Biology 5(2), 12 (2004). doi:10.1186/gb-2004-5-2-r12

49. šåsić, M., šikić, M.: Edlib: a C/C ++ library for fast, exact sequence alignment using edit distance. Bioinformatics 33(9), 1394–1395 (2017). doi:10.1093/bioinformatics/btw753. Publisher: Oxford Academic

50. Köster, J., Rahmann, S.: Snakemake—a scalable bioinformatics workflow engine. Bioinformatics 28(19), 2520–2522 (2012). doi:10.1093/bioinformatics/bts480. Publisher: Oxford Academic

