## supplementary_material for "Enabling multiscale variation analysis with genome graphs"

February 3, 2021

#### Multiscale variation

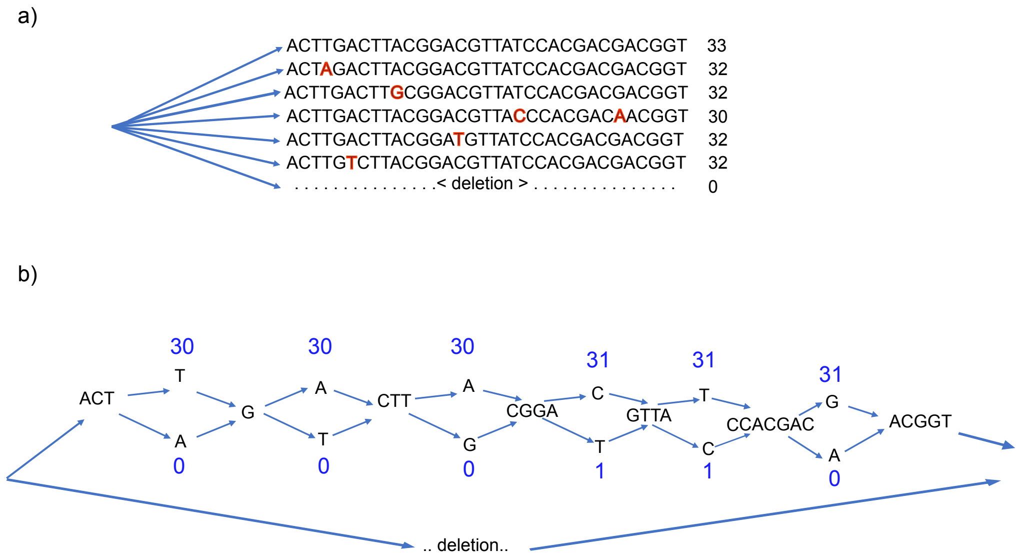

Figure 1: **Comparison of genotyping in non-nested and nested DAGs.** This is intended to show the impact of nesting on genotyping without representing any specific implementation. a) Non-nested graph showing a site with several long alternate alleles differing by SNPs (shown in red). Top allele is the actual allele in a sequenced genome, whose reads have been mapped to this graph. On the right hand end of each allele is the average depth of coverage on this allele. The bottom allele is a deletion, shown with zero coverage. There is very little difference between the true allele and the others which differ by SNPs but only have marginally lower depth of coverage when averaged across the allele. Confidence of the called allele will be low. b) The same sequences in a nested graph, with coverage shown at nested sites. Precise coverage at SNPs makes it very clear what is going on - all SNP calls will be high confidence and correct.

In Fig. 1, we refer to nested and non-nested DAGs; these are formally defined in the Methods section of the paper.

#### Graph constraints

In this section, we illustrate the types of graphs supported by `gramtools`. In Fig. 2, we illustrate the *balanced brackets* property, which graphs must have to be indexed by `gramtools`. As shown in Fig. 2, in some cases, if this property is not satisfied, an alternate graph representation of the same sequences can be produced which does.

In Fig. 3, we illustrate the transformation of a graph with a balanced brackets representation into a graph with the *disjoint subgraphs* property. Graphs with this property are indexed in practice by `gramtools`.

The balanced brackets and disjoint subgraph properties are formally defined in the Methods section of the paper. The Methods section also describes how to produce a balanced brackets representation, and how to transform a graph with balanced brackets into one satisfying the disjoint subgraph property.

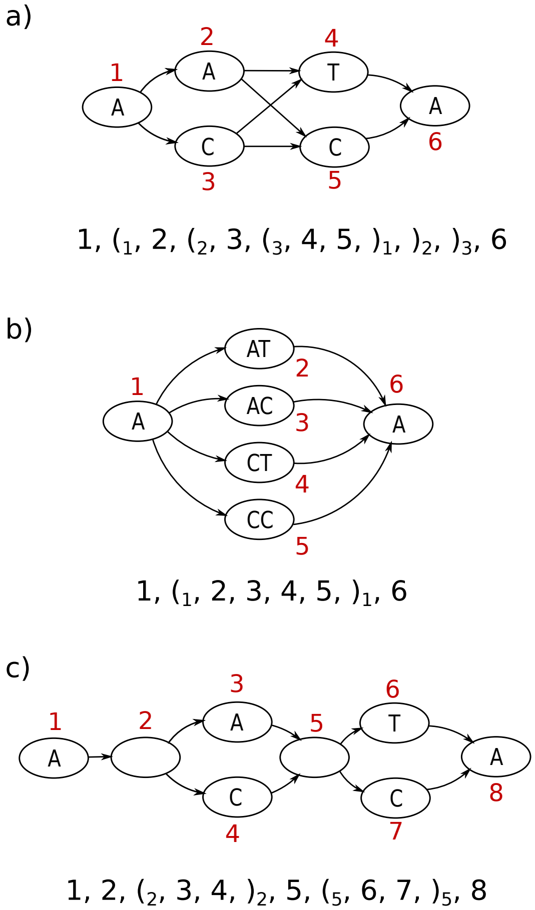

Figure 2: **Indexable and non-indexable graphs representing equivalent sequences.** The graphs in the three panels represent the same set of sequences but have different topology. Each node is labeled with an integer identifier in red, increasing according to a topological ordering of the nodes. The balanced brackets representation of each graph is written below it. The graph in panel a) does not have balanced brackets and cannot be indexed by `gramtools`, while the graphs in panels b) and c) do and can therefore be indexed.

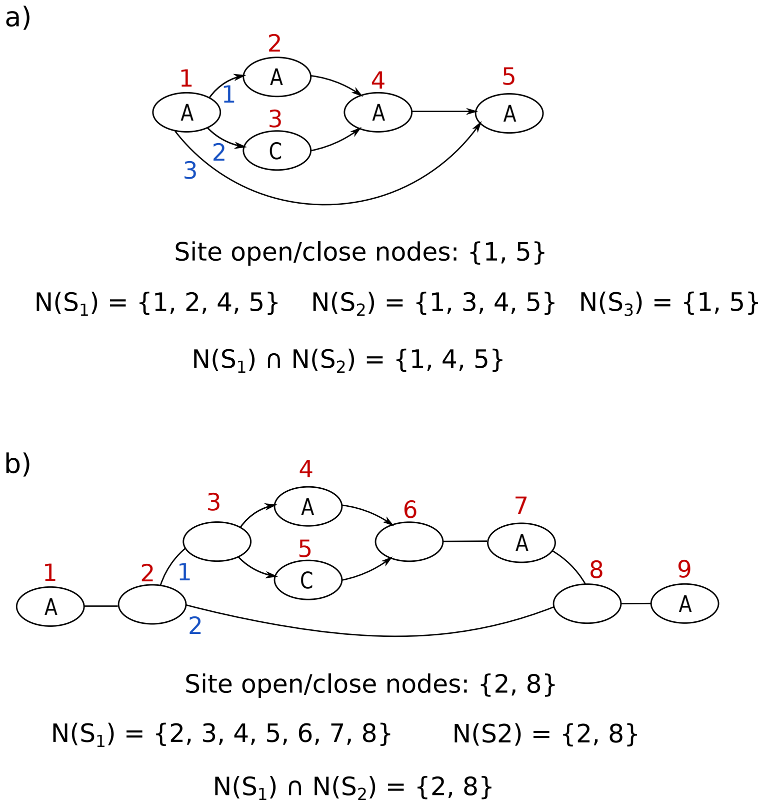

Figure 3: `gramtools` **transformation of indexable graphs.** Each node is labeled with an integer identifier in red, increasing according to a topological ordering of the nodes. In each graph, we focus on one variant site, and the edges outgoing from the site's opening node are labeled in blue. a) This graph satisfies the balanced brackets property (it can be represented as $1, (_1, (_1, 2, 3, )_1, )_1, 5$) but cannot be directly indexed by `gramtools`, as it does not satisfy the disjoint subgraph property. The text below shows why: the intersection of the set of nodes $N(S_1)$ and $N(S_2)$ does not yield only the opening and closing node of the site (namely $\{1, 5\}$). **b)** Once transformed, this graph satisfies the disjoint subgraph property and can be indexed by `gramtools`.

#### JSON output format

The specification of the JSON Variant Call Format (jVCF) produced by `gramtools` is publicly available on the following repository: `https://github.com/iqbal-lab-org/jVCF-spec`.

We provide below the version corresponding to commit 28a955933baadd6c0536856433634c1d14f37ea8 of the repository.

### JSON Variant Call Format Specification

Version 0.1

January 13, 2021

### 1 Rationale

#### 1.1 Variant calls in genome graphs

The Variant Call Format (VCF) is a tab-delimited data format describing genetic variation occurring with respect to a linear reference genome. Here we define an extension to VCF for genome graphs, which are graph structures representing genetic variants occurring on potentially multiple reference genomes.

The format uses JavaScript Object Notation (JSON), a commonly used data format, and we thus call it JSON VCF or jVCF.

#### 1.2 Requirements

jVCF assumes:

- Variant sites have been defined on the genome graph
- Variant sites can be contained in other variant sites. It is known which sites are contained in which others.

For the latter point, a site contained in another occurs in a given sequence background. Each sequence background must be labeled with a unique positive integer, called its **haplogroup**. See this toy graph for an example.

#### 1.3 Description of JSON

The JSON format is defined here: https://www.json.org/json-en.html.

Briefly, JSON consists of *objects* and *arrays*:

- An object is an unordered collection of key/value pairs enclosed in curly brackets ('{' and '}').
- An array is an ordered collection of values enclosed in square brackets ('[' and ']').

In an object, a key is a string. Strings are enclosed in double quotes ('"'). In objects and arrays, values can be:

- a string
- a number
- one of the literals 'true', 'false' or 'null'
- an array
- an object

#### 2 Example

Here is a toy genome graph:

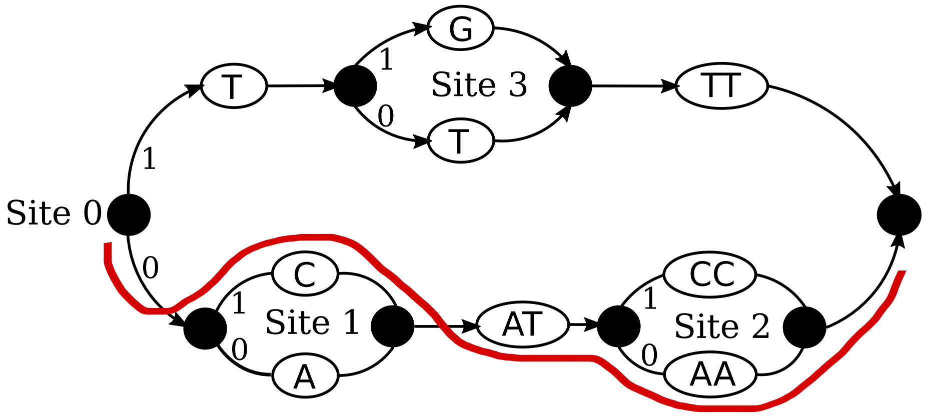

Figure 1: Genome graph with haploid genotyped sample as red path

In Figure 1, black nodes mark site entry and exit points. Each variant site is labeled by a unique positive integer ID, and each outgoing branch from the start of a variant site is labeled by a unique positive integer (its haplogroup).

Assume the ploidy is 1, and we are genotyping a sample whose genotyped path in the graph is the red line. The jVCF of this genotyped sample would look like this:

```
{
  "Sites":[
    {"ALS":["AATAA","CATAA"],"FT":[[]],"GT":[[1]],"HAPG":[[0]],"POS":1,"SEG":"myRef"},
    {"ALS":["A","C"],"FT":[[]],"GT":[[1]],"HAPG":[[1]],"POS":1,"SEG":"myRef"},
    {"ALS":["AA"],"FT":[[]],"GT":[[0]],"HAPG":[[0]],"POS":4,"SEG":"myRef"},
    {"ALS":["T"],"FT":[[]],"GT":[[null]],"HAPG":[[]],"POS":2,"SEG":"myRef"}
  ],
  "Site_Fields": {
    "ALS": {
      "Desc": "Alleles at this site"
    },
    "FT": {
      "Desc": "Filters failed in a sample"
    },
    "GT": {
      "Desc": "Genotype"
    },
    "HAPG": {
      "Desc": "Sample haplogroups of genotyped alleles"
    },
    "POS": {
      "Desc": "Position on reference or pseudo-reference"
    },
    "SEG": {
      "Desc": "Segment ID"
    }
  },
  "Samples": [
    {
      "Desc": "mySample description",
      "Name": "mySample"
    }
  ],
  "Filters": {
    "MINQ": {
      "Desc": "Call is below minimum quality"
    }
  },
  "Model": "myGenotypingModel",
  "Child_Map": {
    "0": {
      "0": [1,2],
      "1": [3]
    }
```

```
  },
  "Lvl1_Sites": [0]
}
```

### 3 Specification

There are 7 required keys in jVCF, which can appear in any order. You can use extra keys beyond the required ones. We list each of the required keys below, giving for each the type and description of its corresponding value.

#### 3.1 Site_Fields

**Type** Object

**Description** Each key describes the fields that can appear in Site objects.

The following keys are required to be present:

| Key | Meaning |
|---|---|
| ALS | Genotyped alleles at this site |
| SEG | Segment (eg chromosome) on which the site lies |
| POS | 1-based position relative to haplogroup reference |
| GT | Genotype calls |
| HAPG | Haplogroup calls: haplogroup the called allele(s) lie on |
| FT | Filters set |

Each key corresponds to an object with at least a “Desc” key which gives a description of the field. Extra keys beyond the required ones can be used.

#### 3.2 Sites

**Type** Array

**Description** Each element is a Site object

##### 3.2.1 Site objects

A Site object describes variant calls at one site. This is analogous to a record in VCF.

As for Site_Fields, the following keys are required to be present:

| Key | Value Type |
|---|---|
| ALS | Array of strings |
| SEG | String |
| POS | Number |
| GT | Array of array of numbers, ‘null’ or empty |
| HAPG | Array of array of numbers or empty |
| FT | Array of arrays of strings or empty |

Here is a more detailed explanation of the keys:

- `ALS`: does not need to store all possible alleles at the variant site. It must store at least one allele which is the reference for this site. The reference allele is the allele obtained by following haplogroup 0 from the start of the site to the end of the site. All called alleles should also be present in `ALS`.
- `SEG`: the name of the genomic segment the site lies on, e.g. chromosome or the name of a reference genome.
- `POS`: 1-based position, offset from the last non-nested site (see Lvl1_Sites). The position of sites contained in other sites is expressed relative to the reference path through the containing site, which is obtained by following haplogroup 0.
- `GT` and `HAPG`: are arrays of arrays. The two array levels are:

  1. Samples. This should have the same size as the array in Samples.
  2. Ploidy. This should have one entry per chromosome copy, or chromosome population (eg tumour subclones or mixed infections).

  For example, to access the first sample's genotype calls at a site, you access the 0th element of the `GT` array. If you have a diploid genotyped sample, the first and second chromosome calls are the 0th and 1st elements of that array.
- `FT`: also an array of arrays. The first level is the samples, and the second the filters set for that sample.

You can use more keys. For each used key, Site_Fields needs to provide a description of its meaning.

##### 3.3 Samples

**Type** Array

**Description** Each entry is an object describing one genotyped sample.

Each entry must contain the following keys: "Name" and "Desc", giving the sample name and its description. The order of the samples matters: it must match that in Site objects.

##### 3.4 Filters

**Type** Object

**Description** Describes the filters that can appear in Site objects.

Each filter that appears under the "FT" key in Site objects must be described here. Each key gives the filter name and each value is an object with at least a "Desc" key giving the filter's description.

#### 3.5 Model

**Type** String

**Description** The name, or a description, of the genotyping model that was used.

#### 3.6 Child_Map

**Type** Object

**Description** Records parent/site relationships between variant sites and on which haplogroups child sites lie.

Each key corresponds to a site index in the Sites array. The value associated to each key is a Child listing object.

Starting from each entry of the Lvl1_Sites, the Child_Map can be recursively traversed to recover the full parent/child site structure of the graph.

##### 3.6.1 Child listing

A Child listing is an object whose keys are haplogroups and whose values are arrays of site indices. It records, for a given parent site, what child sites it has, and which haplogroups the child sites fall under.

#### 3.7 Lvl1_Sites

**Type** Array

**Description** Lists the indices of sites which are not nested in others.

Each element of this array corresponds to a site which has no parents; the value is the index at which to access this site in the Sites array.

Each site referred to in this array may or may not have children, depending on whether or not it appears in Child_Map.

#### Validation of nested genotyping with simulated data

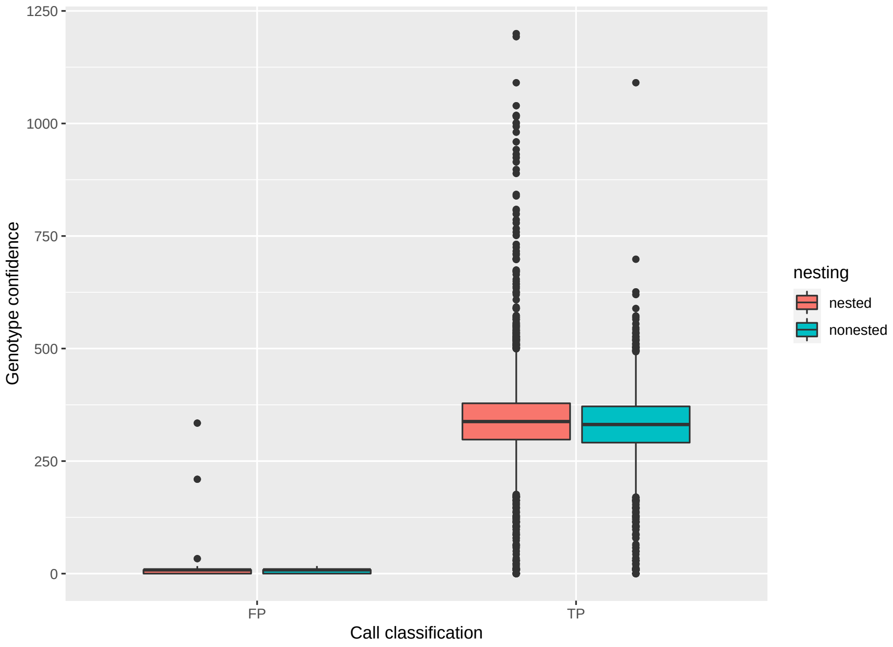

Figure 4: **Improved genotype confidence on nested graphs.** The distribution of genotype confidence (GC) is shown for true positive (TP) and false positive (FP) calls. An FP call counts when the wrong call is made or no call should have been made. Lower overall confidence of FPs allows for filtering.

#### Empirical evaluation against single-reference tools at surface antigens

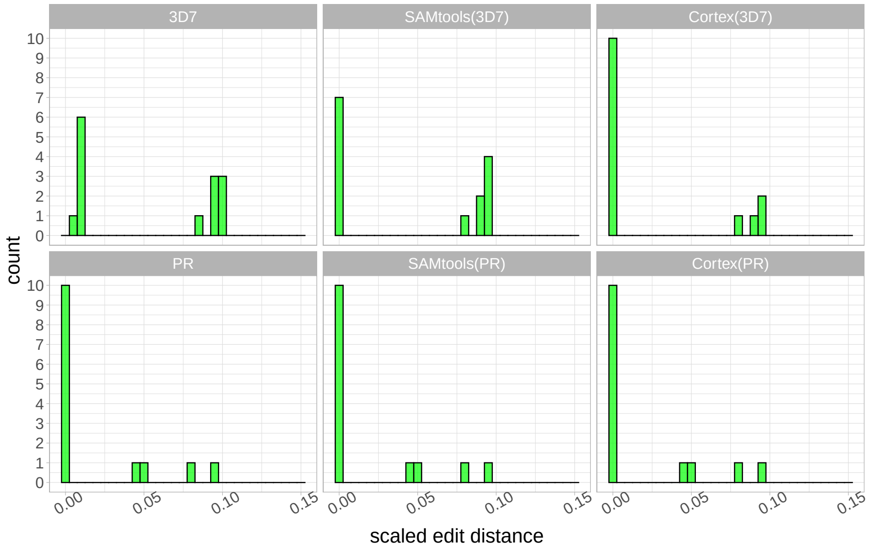

Figure 5: `gramtools` **genotyping compared to reference-based variant calling in DBLMSP.** Four samples are not fully resolved by either `gramtools` or `Cortex` and `SAMtools` run against the personalised reference genome. These samples are candidates for long-read assembly and incorporation into the graph.

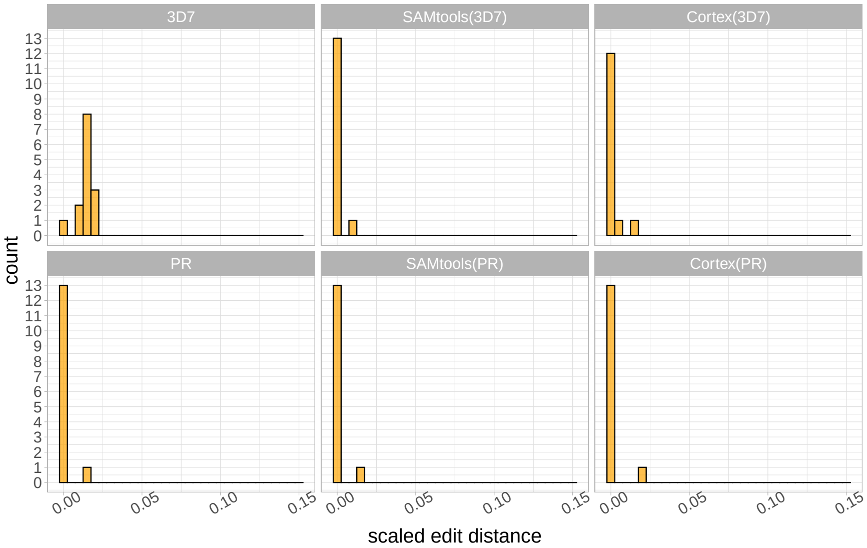

Figure 6: `gramtools` **genotyping compared to reference-based variant calling in AMA1.** All samples except two are perfectly resolved by `gramtools`. The two unresolved samples cannot be improved using `SAMtools` or `Cortex` suggesting, like DBLMSP, long-read assembly is required.

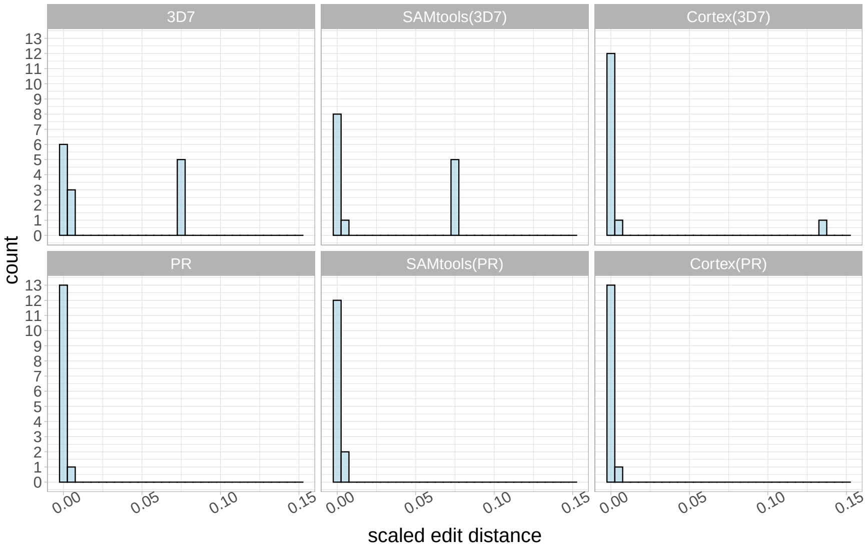

Figure 7: `gramtools` **genotyping compared to reference-based variant calling in EBA175.** All samples are near-perfectly resolved by `gramtools`.

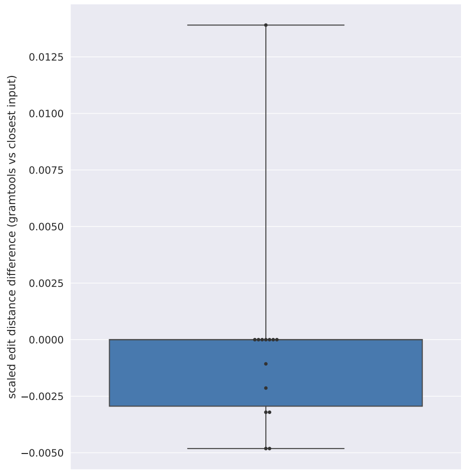

Figure 8: `gramtools`**-inferred sequence compared to the closest possible input in the graph in AMA1.** For all but one sample, the closest input in the graph or a closer recombinant is found.

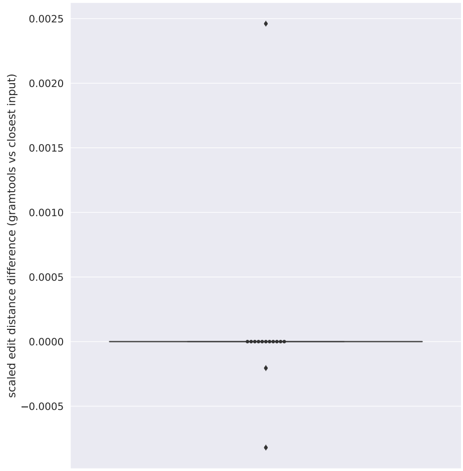

Figure 9: `gramtools`**-inferred sequence compared to the closest possible input in the graph in EBA175.** All but three samples are at a difference of 0, indicating the `gramtools`-inferred gene sequence in those samples is identical to an input haplotype to the graph.

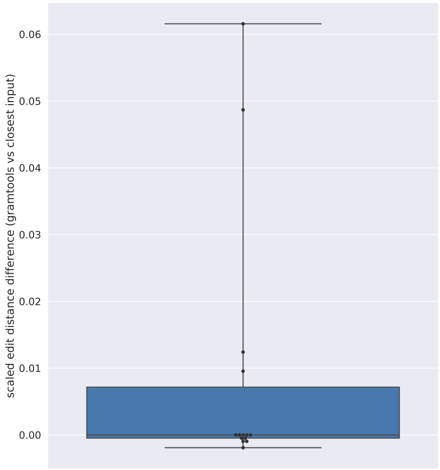

Figure 10: **`gramtools`-inferred sequence compared to the closest possible input in the graph in DBLMSP.** For four samples, `gramtools` did significantly worse than the best possible input sequence in the graph. Investigating the genotyping output revealed a stretch of null calls in a specific region due to no read coverage, causing the reference genome sequence to be called there.

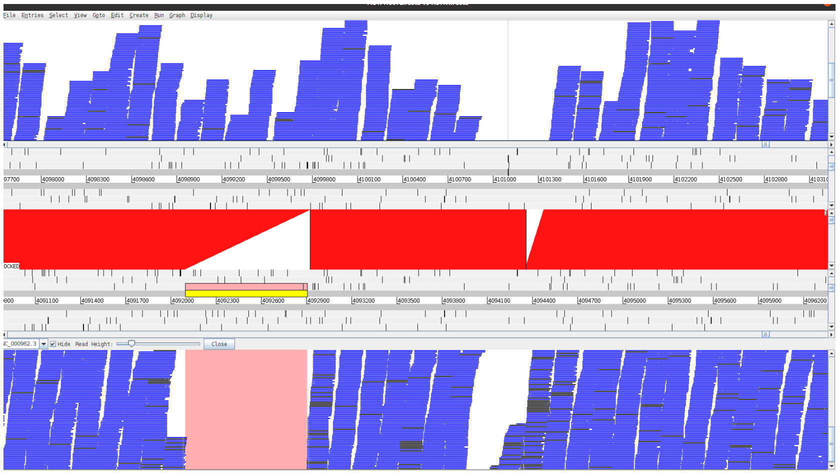

Figure 11: **Manual confirmation of *M. tuberculosis* deletions using `ACT`.** The top track shows the assembly and the bottom track the reference. Mapped reads are shown as blue rectangles. Bright red rectangles show mapped regions between assembly and reference. The light red rectangle highlights the location of the confirmed deletion.

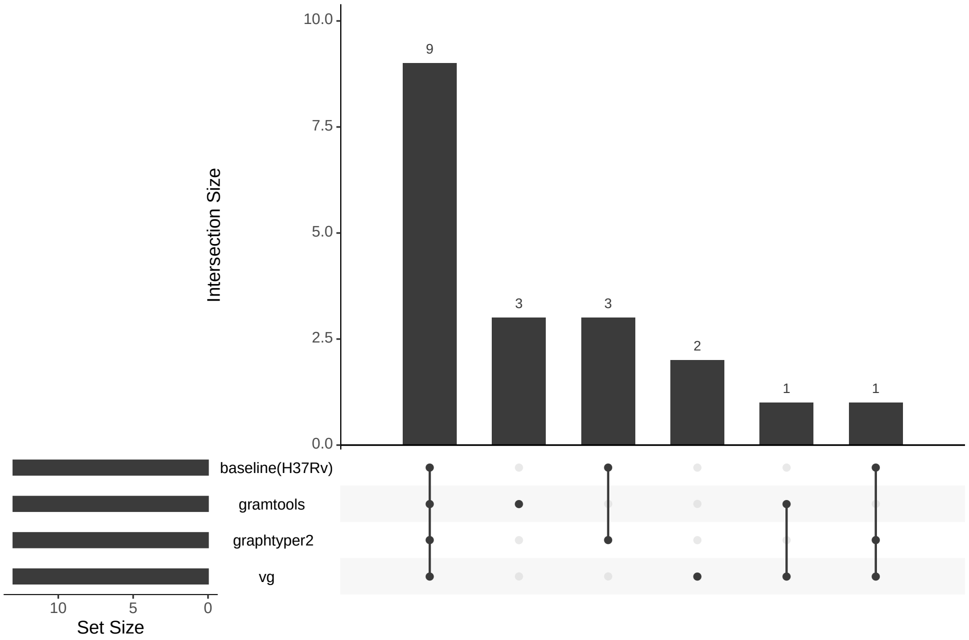

Figure 12: **Unmapped and filtered out sequences in joint SNP and deletion genotyping experiment using `minimap2` based alignments.** Upset plot showing the total number of unmapped sequences and sequences with MAPQ $\leq 30$ and their intersections by condition.

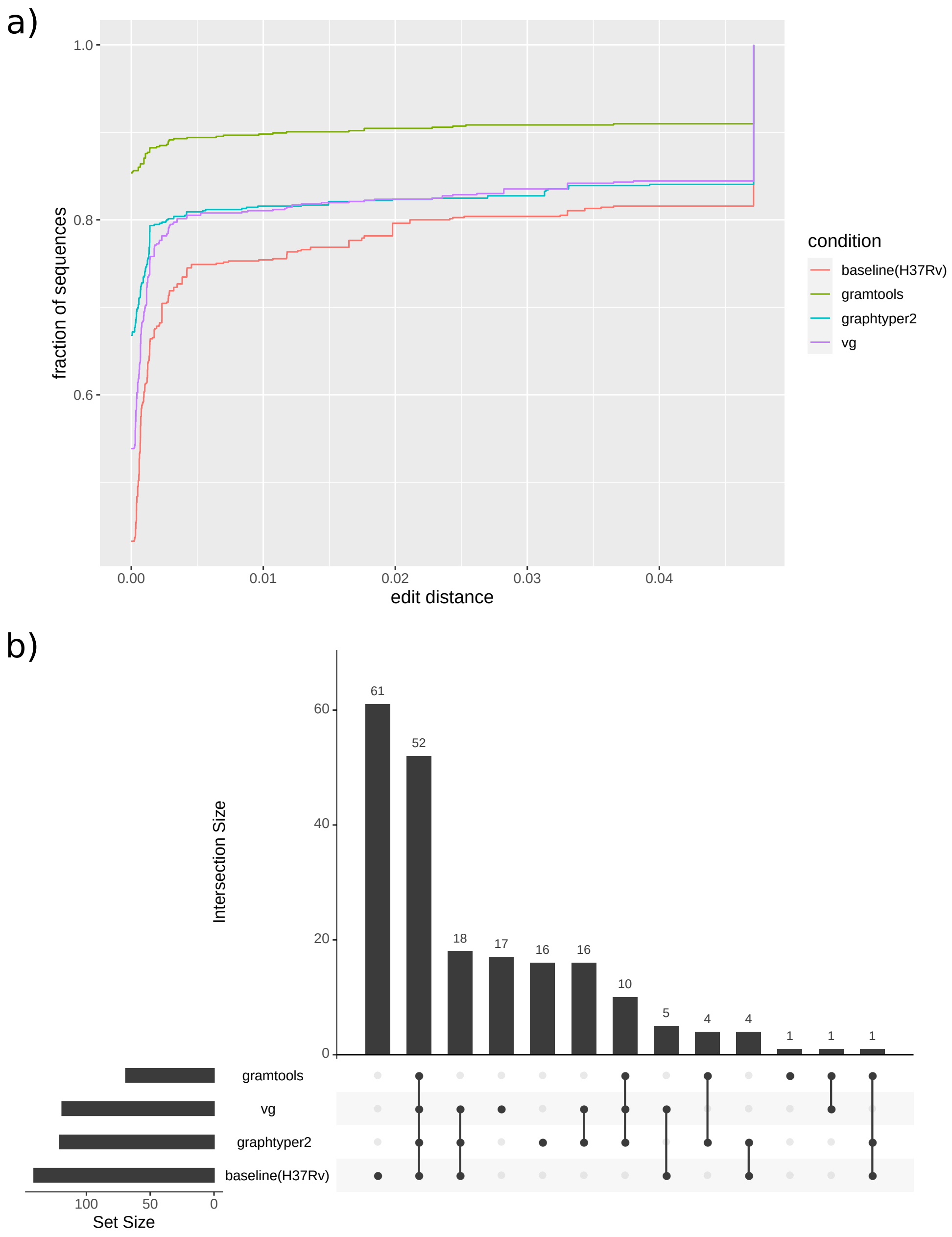

Figure 13: **Joint SNP and deletion genotyping results using `bowtie2` alignments.** a) Cumulative distributions of edit distances between sequences and the truth assemblies. `bowtie2` can only align sequences up to 4.5% divergence making the fraction of aligned sequences lower. Mean edit distance gives the same performance ranking (`gramtools`, `graphtyper2`, `vg`, *baseline*). b) Upset plot of unmapped and MAPQ $\leq$ 30 sequences. For a large number of sequences, the reference genome sequence does not align to the assembly illustrating high divergence between the two.
